# Mitochondrial DNA released by senescent cells triggers immunosuppression in cancer

**DOI:** 10.1101/2023.08.06.551763

**Authors:** Ping Lai, Lei Liu, Nicolò Bancaro, Martina Troiani, Bianca Calì, Jingjing Chen, Prafull Kumar Singh, Rydell Alvarez Arzola, Giuseppe Attanasio, Nicolò Pernigoni, Emiliano Pasquini, Simone Mosole, Andrea Rinaldi, Jacopo Sgrignani, Yuxin Li, Shi Qiu, Pan Song, Yingrui Li, Maria Andrea Desbats, Azucena Rendón Ángel, Ricardo Pereira Mestre, Lucio Barile, Andrea Cavalli, Johann de Bono, Andrea Alimonti

**Affiliations:** Institute of Oncology Research (IOR), Bellinzona, Switzerland; Faculty of Biology and Medicine, University of Lausanne (UNIL), Lausanne, Switzerland; Faculty of Biomedical Sciences, Università della Svizzera Italiana, Lugano, Switzerland; Institute for Research in Biomedicine (IRB), Bellinzona, Switzerland; Department of Urology, Institute of Urology, West China Hospital of Sichuan University, Chengdu, China; Clinical Genetics Unit, Department of Woman and Child Health, University of Padova, Padova, Italy; Laboratory of Cellular and Molecular Cardiology and Laboratory for Cardiovascular Theranostics, Cardiocentro Ticino Foundation, Lugano, Switzerland; Medical Oncology Unit, Institute of Oncology of Southern Switzerland (IOSI), Ente Ospedaliero Cantonale (EOC), Bellinzona, Switzerland; Institute of Cancer Research and Royal Marsden NHS Foundation Trust, London, UK; Department of Medicine, Venetian Institute of Molecular Medicine, University of Padova, Padova, Italy; Department of Health Sciences and Technology (D-HEST), Eidgenössische Technische Hochschule (ETH) Zürich, Zurich, Switzerland

## Abstract

DNA is a potent damage-associated molecular pattern signaling that, once in the extracellular space, triggers the activation of the innate immune system. Here we find that senescent cells release mtDNA to both the cytosol and the extracellular space. In cells undergoing cellular senescence, the release of mtDNA precedes that of nuclear DNA resulting in the activation of the cGAS/STING pathway and establishment of cellular senescence. Intriguingly, by exploiting co-culture and in vivo cross-species experiments, we show that extracellular mtDNA released by senescent tumors cells is specifically captured by polymorphonuclear myeloid-derived suppressor cells (PMN-MDSCs) in the tumor microenvironment (TME). Mechanistically we find that PMN-MDSCs uptake mtDNA to enhance their immunosuppressive ability. Pharmacological inhibition of mtDNA released from senescent tumor cells blocks the PMN-MDSCs immunosuppressive activity, improving the efficacy of therapy-induced senescence (TIS) in cancer. These results reveal the crucial role of mtDNA in initiating cellular senescence and immunosuppression independently of the SASP. Thus, targeting mtDNA release-mediated pathway may hold promise to reprogram the immune suppressive microenvironment in patients treated with chemotherapy.

---

Cellular senescence, a state of stable growth arrest characterized by an intrinsic resistance to apoptosis and a peculiar secretory phenotype termed the senescence-associated secretory phenotype (SASP), has emerged as a critical player in age-related diseases and cancer development^1,2^. Recent studies have shed light on the accumulation of cytosolic nuclear DNA, in the form of cytoplasmic chromatin fragments (CCF), in senescent cells. Once CCF enter the cytoplasm, they are sensed by the GMP-AMP synthase-stimulator of interferon genes (cGAS-STING) pathway that activate the SASP through NF-κB^3–6^. Despite nuclear DNA, cells also contain high copy numbers of mitochondrial circular DNA (mtDNA) densely packaged into nucleoids. Nucleoids include one copy of mtDNA and different proteins, including the mitochondrial transcription factor A (TFAM). In response to various stresses, such as apoptosis^7,8^, viral infection^9,10^, TFAM deficiency^11^, and oxidative stress^12^, mtDNA is released in the cytosol, where it is recognized as a foreign danger to trigger inflammation by the cGAS-STING pathway^11,13,14^. Intriguingly, mtDNA can also be detected in the plasma of patients affected by different age-associated diseases^15,16^, suggesting that stressed cells can also release mtDNA in the extracellular space. A recent report proposes that senescent cells are a primary source of cell-free mtDNA that promote age-associated systemic inflammation^17^.

However, whether and how cytosolic mtDNA release contribute to senescence induction remains elusive. Specifically, the mechanism by which senescent cells release mtDNA outside the mitochondria has never been investigated. Finally, it is unknown whether in TIS, mtDNA release in the extracellular space, promotes immune suppression. Here, we demonstrate that in *Pten* loss-induced cellular senescence (PICS), oncogene-induced senescence (OIS), and therapy-induced senescence (TIS), mtDNA is released outside the mitochondria, through the voltage-dependent anion channel (VDAC). This process requires mitochondrial delocalization of endonuclease G (EndoG). Notably, we also show that in TIS, senescent tumor cells release a high quantity of mtDNA in the extracellular space where mtDNA is selectively internalized by PMN-MDSCs. MtDNA released by senescent tumor cells enhances the immune suppressive and tumor-promoting function of PMN-MDSCs, impacting tumor progression and treatment resistance.

## MtDNA release and senescence induction

To assess whether mtDNA is released in the cytoplasm of cells undergoing different types of senescent response, we used mouse embryonic fibroblasts (MEFs) where senescence was induced by loss of tumor suppressor gene *Pten* (PICS), overexpression of oncogenic *HRasV12* (OIS), and treatment with the CDK4/6 inhibitor Palbociclib (Palbo), a prototypical TIS that does not induce DNA damage^18,19^. In all the cases analyzed, senescent cells exhibited a marked accumulation of cytosolic DNA (indicated by anti-dsDNA antibody) outside the mitochondria (Mitospy) (Fig. 1a, b; Extended Data Fig. 1a-c). We next checked whether the DNA accumulating in the cytosolic fraction of senescence cells was of nuclear or mitochondrial origin by performing qPCR using mitochondrial (Cox1, Dloop) and nuclear (ActB, Tert) specific primers and immunofluorescence staining for DAPI and cGAS to detect CCFs. Cytosolic mtDNA was increased in all the senescent cells analyzed (Fig. 1c). In contrast, cytosolic nDNA was only increased in OIS (Fig. 1c), in line with previous evidence demonstrating an increased nuclear DNA damage and CCF accumulation in this senescent model^3,20^ (Extended Data Fig. 1d, e). Of note, we found that in all the different types of senescent cells, mtDNA was more abundant than nDNA (Fig. 1c). In addition, senescent cells also released more mtDNA than nDNA into the extracellular milieu as detected by qPCR in the conditioned media (c.m) of the different senescent cells (Fig. 1d). Interestingly, released mtDNA of senescent cells was hyper-fragmented, and hypomethylated when compared to control as assessed by Anti-5mC dot blot analysis performed on total mtDNA (Extended Data Fig. 1f, g). We next performed a time course experiment and found that mtDNA release occurred at early time points during senescence establishment in all the senescent models (Fig. 1e). Note that in OIS, where mtDNA and nDNA are increased, mtDNA release preceded that of nDNA (Fig. 1e, f). To understand whether mtDNA is required for the establishment of cellular senescence, we depleted mtDNA in MEFs undergoing PICS, OIS, and TIS through the overexpression of the Herpes Simplex Virus exonuclease UL12.5 that specifically eliminates mtDNA without affecting nDNA^21^. Depletion of mtDNA significantly decreased SA-β-Gal positivity, the p16 and p21 protein levels, and the expression of different SASP-associated genes in all the senescence models (Fig. 1g-j; Extended Data Fig. 1h, i). These data were further validated in TRAMP-C1 murine prostate tumor cells treated with Palbociclib or Docetaxel to induce cellular senescence (Extended Data Fig. 2a-f). Palbociclib induced cytosolic mtDNA release, Docetaxel treatment increased the release of both mtDNA and nDNA. However, docetaxel treated TRAMP-C1 cells released higher amount of mtDNA than nDNA in the cytosol and the culture media (Extended Data Fig. 2c, d). Moreover, depletion of mtDNA in TRAMPC1 cells infected with UL12.5 and treated with Palbociclib abrogated senescence induction and SASP genes expression (Extended Data Fig. 2g-j). Together, these data highlight the crucial role of mtDNA in senescence in both primary MEFs and cancer cells.

**Fig. 1.**
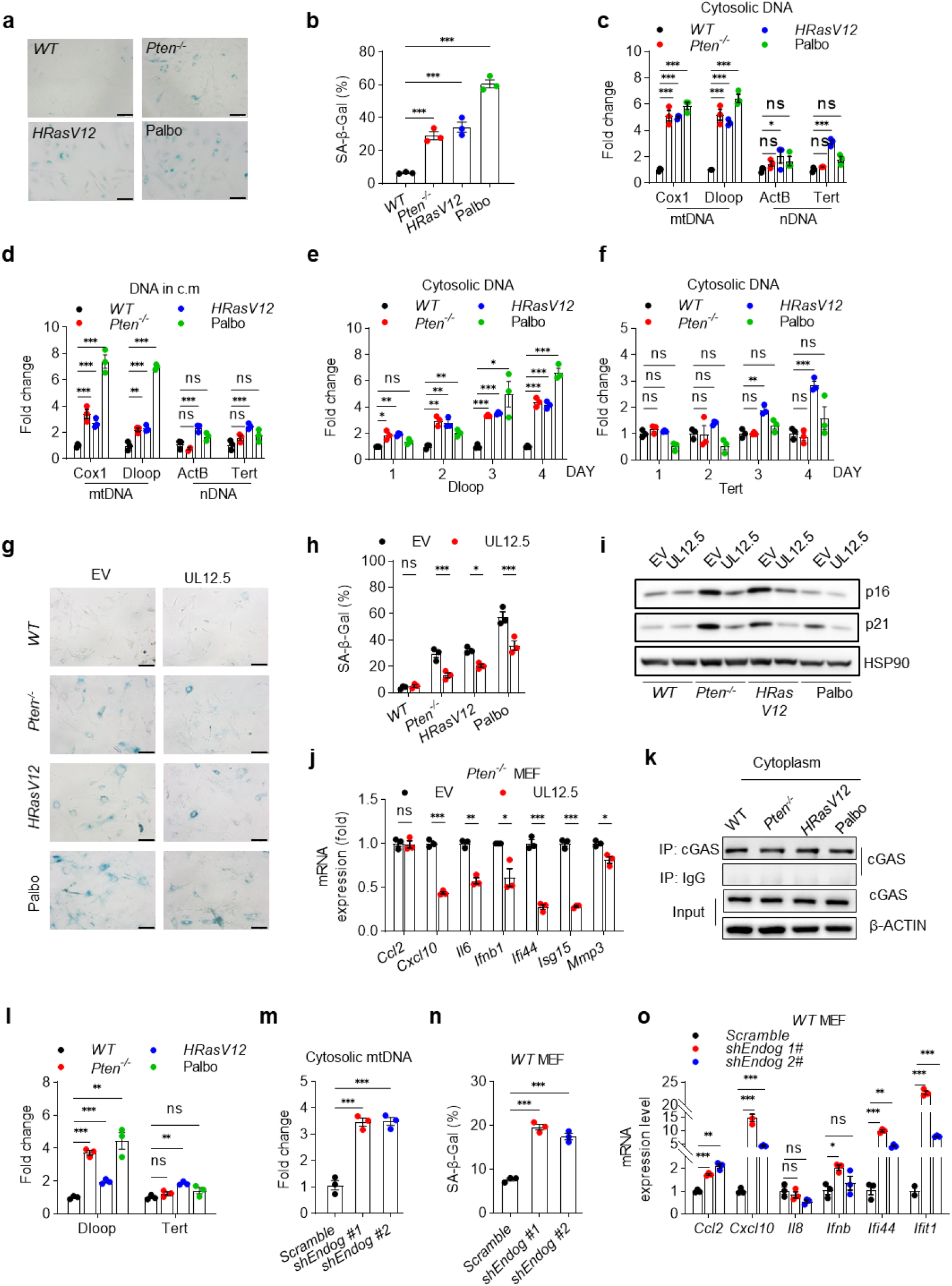
mtDNA release is required for senescence induction. **a,b,** Representative images of SA-β-gal of three different senescent types (scale bar: 200 µm) (**a**) with quantification in the percentage of SA-β-gal positive cells (**b**), n = 3 per group. **c,** Quantification of cytosolic mtDNA and nDNA in senescent MEFs. **d,** Quantification of mtDNA and nDNA in conditioned medium (c.m) collected from indicated senescent MEFs. **e, f,** Quantification of cytosolic Dloop (mtDNA) **(e)** and Tert (nDNA) **(f)** change in different time points during senescence induction. **g-j**, MEFs were transduced with HSV1 UL12.5 or empty vector (EV), and senescence was induced as aforementioned. SA-β-gal staining and quantification (**g, h**), protein (**i**), and SASP gene expression (**j**) were examined in indicated models. **k**, Cytoplasm of senescent MEFs were immunoprecipitated with anti-cGAS antibody or rabbit IgG. Immunoblot analysis of cGAS precipitation was shown. **l**, Precipitated DNA was amplified by real-time qPCR using the indicated primer pairs. **m**, Quantification of cytosolic mtDNA in *WT* MEFs infected with *shEndog* lentivirus. **n,** SA-β-gal in *WT* MEFs after *Endog* knocking down. **o**, Indicated SASP genes expression were detected in *WT* MEFs after *Endog* knocking down. Data pooled from one experiment representative of at least three independent experiments (**a, c-g, j, l, m, o**) or three independent experiments (**b, h, n**) are shown (mean ± SEM). Two-way ANOVA followed by Dunnett’s multiple comparisons test was used to evaluate the statistical significance in **h**, One-way ANOVA followed by Tukey’s multiple comparisons test was used in **b-f, l-o**. Multiple unpaired t-test was used in **j**. *p < 0.05; **p < 0.01; ***p < 0.001; ns, not significant.

We next hypothesized that increased mtDNA release in senescent cells could activate the cGAS-STING pathway as previously reported for CCF in OIS^3,22^. Immunoprecipitation of cGAS in the cytoplasmic fraction of control and senescent cells showed an increased binding of mtDNA rather than nDNA to cGAS in PICS and TIS (Fig. 1k, l). Whereas, in OIS, we detected both mtDNA and nDNA bound to cGAS, in line with previous evidence^3^ (Fig.1l). Knockdown of cGAS (via short-hairpin *shMb21D1*) in MEFs infected with Cre, HRasV12, or treated with Palbo reduced p21, p27 protein levels, and the expression of SASP genes in all the models analyzed (Extended Data Fig. 3a-d). To finally assess whether cytosolic mtDNA is required for senescence induction independently of cytosolic genomic DNA, we inactivated Endonuclease G in wild-type (*WT*) MEFs using shRNAs (Extended Data Fig. 3e). EndoG is the most abundant and active mitochondrial endonuclease, located in the intermembrane space of mitochondria, and it is involved in the degradation of damaged mtDNA^23,24^. Previous evidence demonstrates that cells depleted of EndoG release high levels of mtDNA outside the mitochondria^25^. Notably, EndoG downregulation in *WT* MEFs increased the amount of cytosolic mtDNA, induced cellular senescence, and promoted the upregulation of SASP genes (Fig. 1m-o; Extended data Fig. 3f). Together these findings demonstrate that cytosolic mtDNA release is required for senescence induction and activation of the SASP.

## ROS and EndoG regulate mtDNA release in senescent cells

Reactive oxygen species (ROS) accumulation is associated with mitochondrial dysfunctions in senescent cells^20,26–29^. Since mtDNA release was found in all the senescent models, we speculated that increased ROS production in mitochondria may cause the release of mtDNA. The staining of CM-H2DCFDA, a fluorescent indicator for ROS, showed that ROS production was increased in all the senescence models analyzed (Extended data Fig. 4a). Moreover, the depletion of mitochondrial ROS by mito-TEMPO, a mitochondria-targeted antioxidant, prevented cytosolic mtDNA accumulation as well as cellular senescence and SASP-associated genes in PICS and OIS (Extended data Fig. 4b-h). In line with these findings, oxidative stress induced by low dosage H_2_O_2_ was sufficient to cause the cytosolic mtDNA release in *WT* MEFs and induced cellular senescence (Extended data Fig. 4i, j). Together, these data indicate that ROS facilitates cytosolic mtDNA release to induce cellular senescence and SASP production.

In physiological conditions, the cellular pool of mtDNA is maintained by mitochondrial endonuclease, whose mitochondrial to nuclear localization is regulated by ROS^30,31^. Having demonstrated that ROS elevation and EndoG inactivation are required for the cytosolic release of mtDNA, we assessed the levels and location of EndoG in PICS, OIS, and TIS. While we did not detect any changes in EndoG levels in senescent vs. control cells (Extended data Fig. 5a), immunofluorescence images and immunoblot with different cell fractions showed that in senescent cells, EndoG translocated from the mitochondria to the nucleus (Extended data Fig. 5b-d). During apoptosis, nuclear EndoG localization promotes chromosomal DNA fragmentation via interaction with apoptosis-inducing factor (AIF)^32–34^. However, AIF did not translocate into the nucleus with EndoG in senescent cells (Extended data Fig. 5d, e), in line with the known notion that senescent cells are resistant to apoptosis^35^. Together, these results suggest that ROS induces mtDNA release and EndoG translocation in senescent cells, thereby protecting mtDNA from degradation by this endonuclease.

## Senescent cells release mtDNA through VDAC

We next focused on the mechanism by which mtDNA is released outside the mitochondria. In apoptosis, mtDNA release is mediated by macropores in the mitochondrial outer membrane (MOM) created by oligomerization of the proteins BAX and BAK^9^. In cells subjected to oxidative stress, mtDNA can escape through macropores formed by mitochondrial permeability transition pores (mPTP) and oligomerizing voltage-dependent anion channels (VDAC1)^7,25^. We therefore checked whether mtDNA release from senescent cells could be blocked by treatment with a BAX inhibitor (BAX Inhibiting Peptide V5, BIP-V5) and mPTP inhibitor (cyclosporin A, CsA). However, qPCR analysis performed in the cytosolic fraction of senescent cells showed no decrease in mtDNA release upon treatment with these compounds (Extended data Fig. 6a). We then investigated whether VDACs could mediate the cytosolic mtDNA release in senescent cells. Knocking down *Vdac1* or *Vdac2* reduced the cytosolic release of mtDNA and senescence in PICS (Extended data Fig. 6b-e). In addition, cells undergoing senescence treated by VBIT-4, a selective inhibitor of VDAC1 oligomerization, impaired the release of mtDNA in the cytosol with consequent interference in the establishment of cellular senescence and SASP production in both senescent MEFs and cancer cells treaded with TIS (Extended data Fig. 6f-o). Taken together, our findings demonstrate that VDACs are involved in the release of mtDNA during cellular senescence and that pharmacological inhibition of VDAC in senescent cells may work as a therapeutic strategy to suppress the detrimental effect of the SASP in TIS^36^.

## PMN-MDSCs uptake mtDNA from senescent tumor cells

Senescent cells have been shown to exert detrimental effects on the tissue microenvironment, contributing to tumorigenesis through different mechanisms^37,38^. In prostate cancer, we and others have shown that senescent tumor cells enhance the recruitment of different populations of myeloid cells^1,39,40^. As demonstrated, PMN-MDSCs are the main myeloid immune subset infiltrating both mouse and human prostate cancers, whereas T and NK cells are scarce^1,39^. To better characterize the role of mtDNA released by senescent tumor cells in the tumor microenvironment, we assessed whether extracellular mtDNA could be uptaken by myeloid cells through a trans-species experiment using human prostate tumor cells injected in NRG mice. To do so, we first evaluated whether also human senescent tumor cells could release mtDNA equally to senescent MEFs and mouse tumor cells. PC3, prostate tumor cells were treated with Palbociclib and Docetaxel to induce cellular senescence (Fig. 2a, b). Senescence induction by drug treatments was accompanied by the release of mtDNA in both the cytosol and the extracellular space (Fig. 2c, d). We next injected PC3 cells in NRG mice and treated them with or without Palbociclib to induce senescence *in vivo*. Different populations of murine myeloid cells were then sorted from tumors of untreated and treated mice by using the following markers: CD11b^+^CD11c^+^ (dendritic cells), CD11b^+^F4/80^+^ (macrophages) and CD11b^+^ Ly6G^+^ (PMN-MDSCs). Finally, qPCR analysis was performed in the different immune subsets to detect mtDNA using human-specific primers (Fig. 2e and Extended Data Fig. 7a). We found that in mice treated with TIS, tumor-infiltrating CD11b^+^ Ly6G^+^ cells accumulate a significantly higher amount of human mtDNA (Nd1) compared to other sorted immune populations (Fig. 2f). Of note, no change of human nuclear DNA (LINE1) was detected in the sorted immune cell populations thereby validating our previous findings in the c.m of cells treated with Palbo (Fig. 2f).

**Fig. 2.**
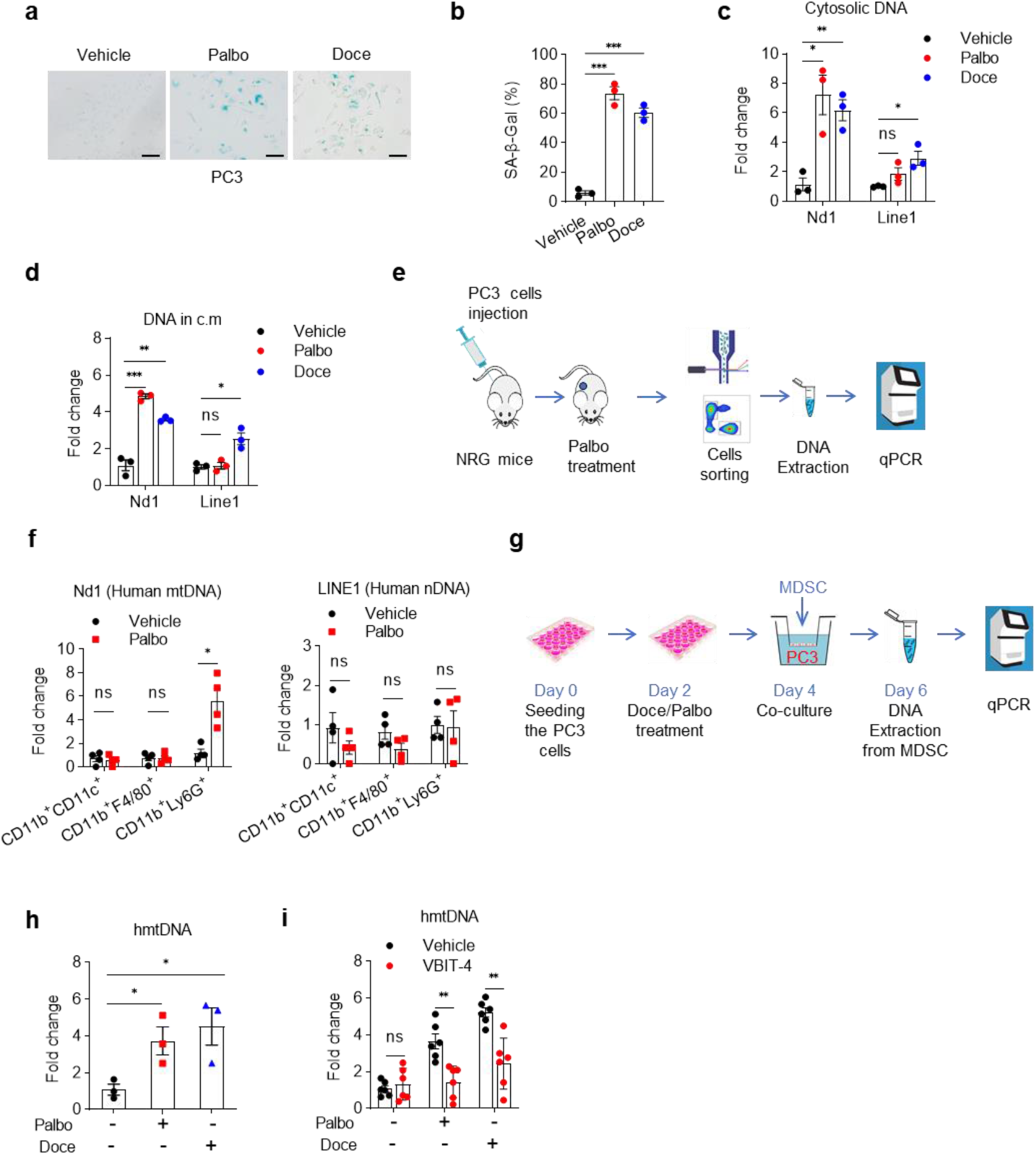
mtDNA release from senescent tumor cells in the extracellular space is uptaken by PMN-MDSCs. **a**, **b**, Representative images (**a**) and quantification (**b**) of SA-β-gal of PC3 cells treated with 2.5 µM Palbociclib (Palbo) or 10 nM Docetaxel (Doce) for 48 h. **c, d**, Quantification of mtDNA and nDNA release in the cytosol (**c**) and c.m (**d**) in TRAMP-C1 treated with Palbo and Doce as mentioned before. **e**, Scheme of experimental design. **f**, Fold change of human mtDNA and nDNA that originated from PC3 tumor cells were quantitated by q-PCR using specific primers and normalized to mouse total DNA which, obtained from the whole-cell extract of mouse CD11b^+^ CD11c^+^, CD11b^+^ F4/80^+^ and CD11b^+^ Ly6G^+^ myeloid cells in the tumor. **g**, Scheme of timeline and experimental design of human PC3 cell co-cultured with mouse BM-MDSC. **h**, qPCR analysis of human mtDNA in mouse BM-MDSC co-cultured with senescent PC3 cells induced with Palbo and Doce. **i**, qPCR analysis of human mtDNA in mouse BM-MDSC co-cultured with PC3 cells treated with Palbo or Doce combined w/o VBIT-4. Data pooled from one experiment representative of at least three independent experiments (**c, d, h, i**) or multiple independent experiments are shown (**b**), each dot represents the value from one mouse in **f** (mean ± SEM). One-way ANOVA followed by Tukey’s multiple comparisons test was used in **b-d**, **h**. Multiple unpaired t-test was used in **f** and **i**. *p < 0.05; **p < 0.01; ***p < 0.001; ns, not significant.

To further validate these findings in vitro, we co-cultured bone marrow-derived murine MDSCs (BM-MDSCs) with senescent human PC3 cells. PC3 cells were pre-treated with Palbo and Doce to induce cellular senescence. Upon senescence establishment, cells were incubated with BM-MDSCs for 48h. qPCR analysis in murine MDSCs showed an increased accumulation of human mtDNA (Fig. 2g, h). These data were validated using murine TRAMP-C1 and MDSCs cells. To perform this experiment, murine TRAMPC-1 were pre-incubated with BrdU to label the DNA before treatment with Palbo and Doce. FACS analysis for BrdU in MDSCs showed the presence of tumor cells derived DNA (Extended data Fig. 7b-d). Finally, treatment with the VDAC1 inhibitor VBIT-4 in senescent human prostate tumor cells decreased the cytosolic and extracellular levels of mtDNA and the mtDNA uptake by murine BM-MDSC (Fig. 2i). Collectively, these results suggest that PMN-MDSCs are capable of internalizing extracellular mtDNA that originates from both human and murine senescent tumor cells.

## MtDNA released by senescent tumor cells enhances the immune suppressive function of MDSCs though activating the cGAS signaling pathway

Having demonstrated that the MDSCs internalize mtDNA derived from senescent tumor cells, we investigated whether senescent cells could affect MDSCs function through the extracellular release of mtDNA. To address this point, we collected c.m from senescent tumor cells and performed DNase digestion to eliminate mtDNA contained in the c.m (Fig. 3a; Extended data Fig. 7e). DNA undigested and digested c.m. was later used to treat BM-MDSCs and perform an immunosuppressive assay using splenic T cells. BM-MDSCs treated with c.m from senescent cells showed an increased immunosuppressive capability in comparison to c.m from non-senescent cells (Pablo vs. Vehicle) (Fig. 3a). However, this effect was decreased by treatment with DNase that eliminated mtDNA from the c.m thereby demonstrating that mtDNA has a direct role on promoting MDSCs activation which is independent of the SASP (Fig. 3b). To validate these findings, we next directly transfected mtDNA extracted from c.m of senescent cells in BM-MDSCs. Transfection of mtDNA significantly increased the expression of pro-inflammatory genes in BM-MDSCs and enhanced their T cells immunosuppressive ability (Fig. 3c, d). RNA-seq analysis of BM-MDSCs transfected with mtDNA provided evidence that mtDNA exhibited distinct abilities to upregulate pro-inflammatory genes, which were strongly associated with immunosuppressive signatures of MDSCs (Fig. 3e, f). Stimulation of MDSCs with mtDNA also significantly changed the expression levels of several genes involved in immune suppression and tumor promotion, such as IL6 signaling^41,42^ (Extended data Fig. 7f, g).

**Fig. 3.**
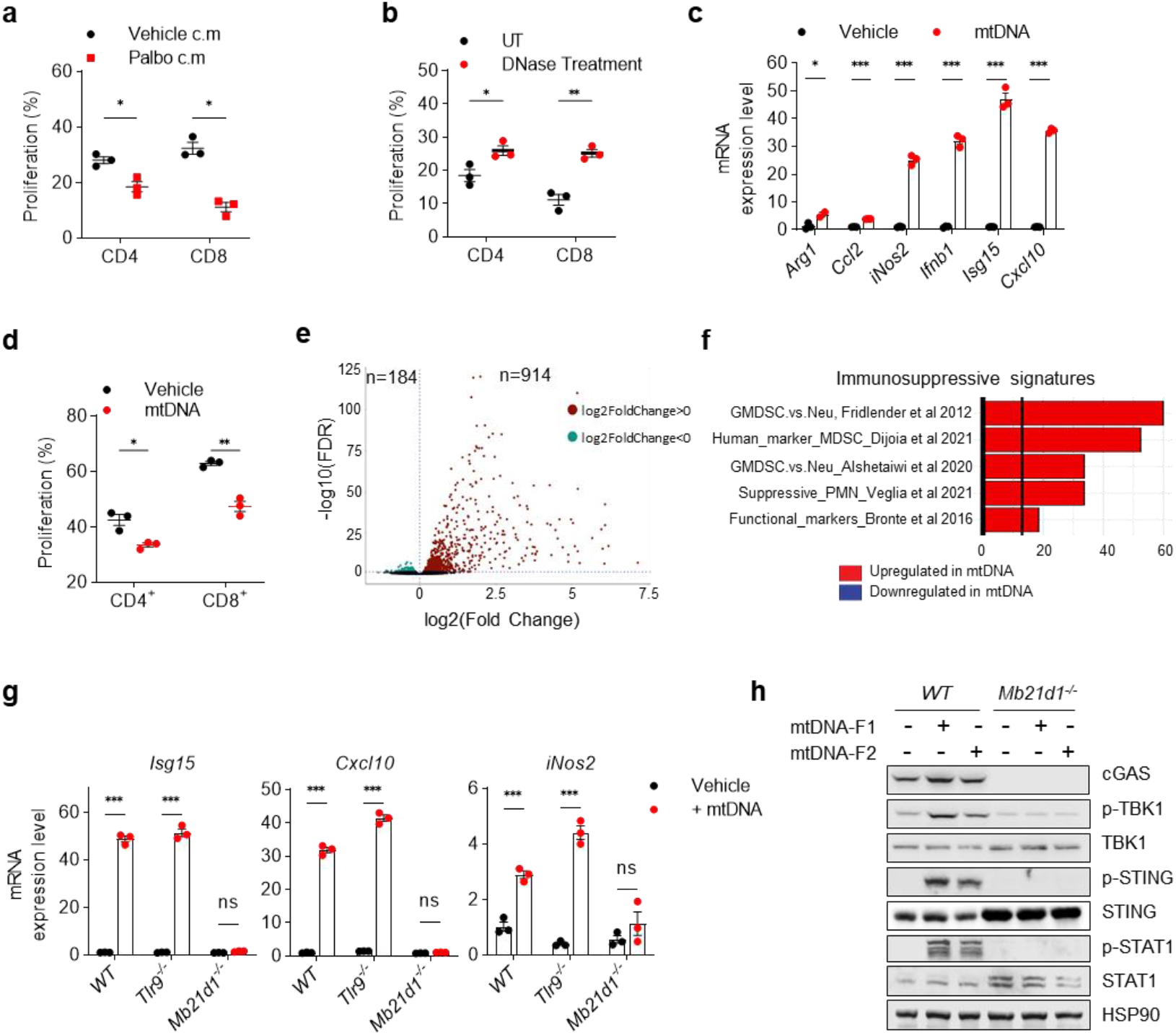
Senescent cells-derived mtDNA increases the immune suppressive function of MDSCs. **a**, Suppression assay of T cell proliferation by BM-MDSC pre-treated with c.m collected from Palbo treated and untreated TRAMP-C1 for 24 h. BM-MDSC and splenocytes were seeded at a ratio of 1:10. **b**, c.m from Palbo-treated TRAMP-C1 cells were performed DNase digestion. Suppression assay of T cell proliferation by BM-MDSC pre-treated with different c.m. **c,** BM-MDSC were transfected with mtDNA at a concentration of 200 ng/ml for 12 h. mRNA expression levels of the indicated genes were detected by RT-qPCR. **d**, Quantification of suppression assay of T cell proliferation by BM-MDSC pre-transfected with mtDNA as mentioned before. BM-MDSC and splenocytes were seeded at a ratio of 1:10. **e**, Volcano plot of RNA-seq of BM-MDSC transfected with or without mtDNA. The dots indicate the differentially expressed genes with Log2FoldChange>0 and FDR<0.05. **f**, Immunosuppressive signatures analysis of upregulated genes in previous samples. **g**, RT-qPCR analysis of the indicated genes expression level in *WT*, *Tlr9^-/-^*and *Mb21d1^-/-^* BM-MDSC after mtDNA transfection. **h**, WB analysis of cGAS-STING signaling pathway activation in *WT* and *Mb21d1^-/-^* BM-MDSC transfected with two different fragmented mtDNA. Data pooled from one experiment representative of at least three independent experiments (**a-d**, and **i**) are shown (mean ± SEM). Multiple unpaired t-test was used in **a-d**, **g**. *p < 0.05; **p < 0.01; ***p < 0.001; ns, not significant. In **e**, FDR<0.05.

We next assess the mechanism by which mtDNA enhances the immune suppressive activity of MDSCs by transfecting mtDNA in BM-MDSCs wild type or knock out for the TLR9 (*Tlr9*^-/-^) or cGAS (*Mb21d1*^-/-^). We found that mtDNA failed to activate MDSCs function in cGAS knock-out cells. In contrast, deficiency of TLR9 receptor in BM-MDSCs was ineffective (Fig. 3h). Western blot analysis in BM-MDSCs transfected with two different mtDNA fragments showed that mtDNA increased the phosphorylation of TBK1, STING, and STAT1 (Fig. 3i). However, upregulation of pSTING, pTBK1 and pSTAT1 was abolished in cGAS knock-out BM-MDSCs. These results indicate that mtDNA-induced pro-inflammatory genes expression in BM-MDSC depends on the activation of the cGAS-STING pathway.

## Blockage of mtDNA release improves TIS efficacy

We next assessed whether blockage of extracellular mtDNA release in senescent cells could enhance the efficacy of Palbociclib treatment. Since VBIT4 treatment decreased the levels of mtDNA release in the cytosol and extracellular space of senescent cells, we hypothesized that this compound could work as both a senostatic by reducing the SASP and direct anti-inflammatory agent by affecting the activation of PMN-MDSCs in tumors treated with Palbociclib. First, we treated TRAMP-C1 tumor-bearing mice with Palbociclib to induce senescence; next, we administered VBIT-4 to prevent the release of mtDNA and thereby inhibit the SASP and the simultaneous activation of PMN-MDSCs.

We found that VBIT-4 increased the efficacy of Palbociclib treatment by decreasing tumor size and frequency of intratumoral CD11b^+^Ly6G^+^Ly6C^int^ cells without affecting the number of these immune cell populations in the blood, spleen, and lymph node (Fig. 4a, b, Extended data Fig. 8a-c). Of note, Palbociclib increased the levels of mtDNA in the plasma of treated mice (Fig. 4c). However, VBIT-4 decreased the level of plasma mtDNA of Palbo-treated mice without affecting the levels of nDNA (Fig. 4c, d). These data were further validated in the RapidCap (PTEN/p53-deficient) allograft prostate tumor model where Palbociclib was administered alone or in combination with VBIT-4 (Extended data Fig. 8d-f). In PMN-MDSCs sorted from tumors treated with Palbo and VIBIT4, we detected a decreased expression of *iNOS2* and *S100A8* in line with the decreased immunosuppressive activity of these cells (Fig. 4e, f). FACS analysis performed in these cells also showed reduced activation of the cGAS-STING pathway in MDSCS from mice treated with Palbo and VIBIT4 compared to Palbo-treated mice (Fig. 4g, Extended data Fig. 8g).

**Fig. 4.**
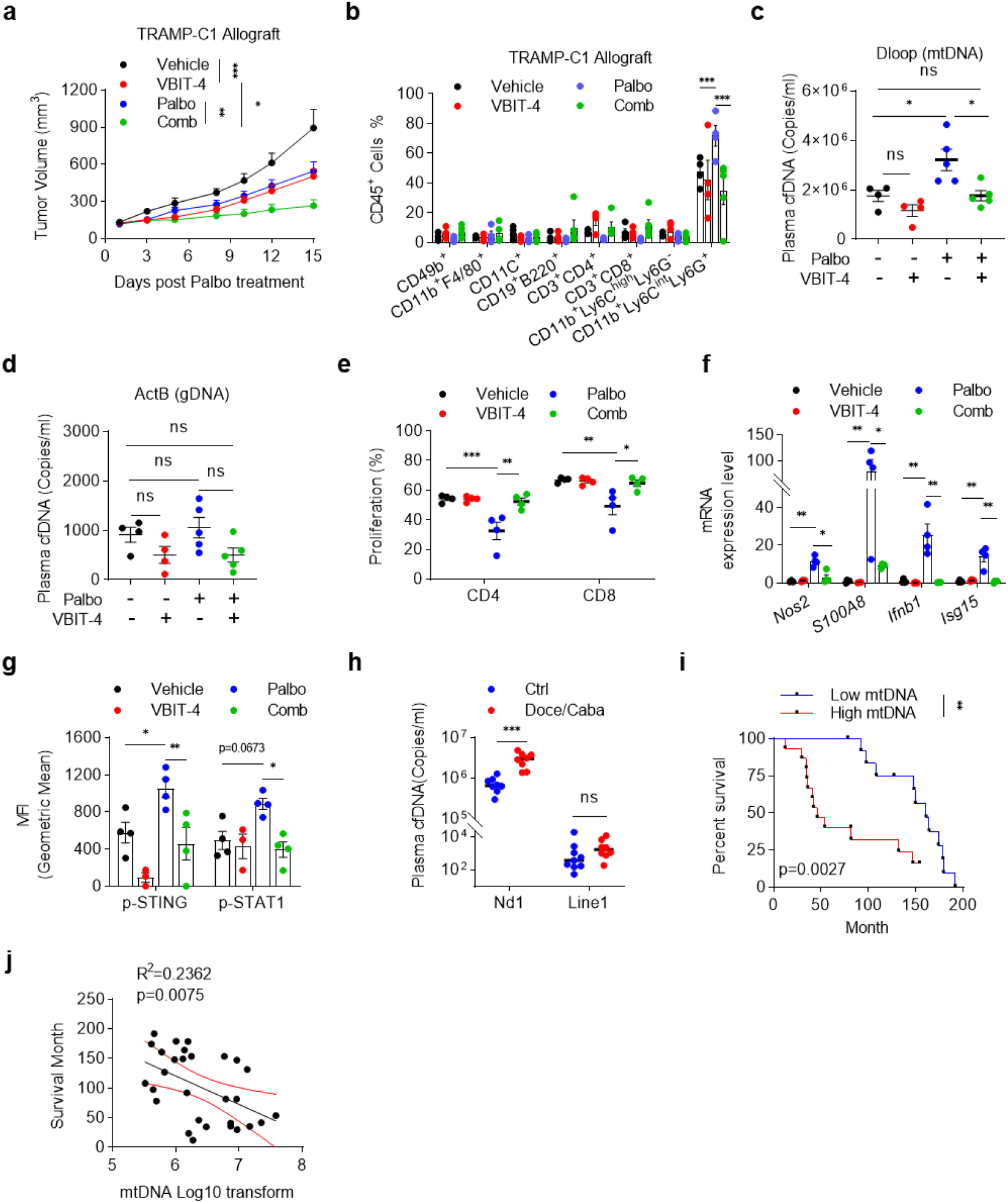
VBIT-4 treatment blocks mtDNA release enhancing the efficacy of TIS. **a**, TRAMP-C1 tumor growth in mice with Palbo, VBIT-4 or combination treatment. UT: n =6, VBIT-4: n=5, Palbo: n=9, Comb: n=5. **b**, Percentages of tumor-infiltrating immune cell populations (gated on CD45^+^ cells). **c**, **d**, Absolute quantification of plasma cell-free mtDNA (**c**) and nDNA (**d**) copy number in TRAMP-C1 tumor-bearing mice with Palbo, VBIT-4, or combination treatment. **e**, Immunosuppressive activity by CD11b^+^ Ly6C^int^ Ly6G^+^ PMN-MDSC sorted from TRAMP-C1 tumors as mentioned before. **f**, RT-qPCR analysis of indicated genes in sorted PMN-MDSC as before. **g**, Fluorescence intensity of p-STING1 and p-STAT1 in CD11b^+^ Ly6c^int^ Ly6G^+^ PMN-MDSC from TRAMP-C1 tumors as before. **h**, Absolute quantification of plasma cell-free nDNA and mtDNA copy number in prostate cancer patients w/o Docetaxel or Cabazitaxel treatment., **i**, Survival curves of prostate cancer patients with low versus high plasma mtDNA copy numbers. MtDNA copy numbers of 28 patients’ plasma were measured by absolute qPCR. According to the median value of mtDNA copy number, patients were equally divided into two groups (High mtDNA> 2285524 copies/ml, Low mtDNA<2285524 copies/ml). **j**, Pearson’s correlation between plasma mtDNA copy number and patients’ survival in months. Data pooled from each dot represent the value from one mouse or patient (**b-h**, **j**). Two-way ANOVA followed by Dunnett’s multiple comparisons test was used to evaluate the statistical significance in **a-g**. Multiple unpaired t-test was used in **h**. For **j**, *p* values were calculated between the High and Low mtDNA groups using log-rank (Mantel-Cox) tests. *p < 0.05; **p < 0.01; ***p < 0.001; ns, not significant.

Clinically, we found that patients treated with Docetaxel or Cabazitaxel, two compounds that drive senescence in cancer cells and normal tissues, exhibited a significant increase of cell-free mtDNA in the plasma, supporting our finding that TIS-induced mtDNA release (Fig. 4h). Finally, we found that patients with high mtDNA circulating levels had significantly lower survival than patients with low circulating mtDNA levels (Fig. 4i, j). These data demonstrate that the blockade of mtDNA release in TIS during prostate cancer therapy can reprogram tumor-infiltrating PMN-MDSCs, thereby promoting the antitumor effects.

## Discussion

DNA localization in the cytoplasm of a cell, whether exogenous or endogenous, acts as a potent danger signal that stimulates an innate immune response. Previous research has demonstrated that CCF^3^ and genomic retrotransposable elements (TE) sequences LINE-1 and endogenous retrovirus (ERV) cDNA^4,5,43^ could accumulate in the cytoplasm and activate cytoplasmic pro-inflammatory pathways in senescence and cancer. However, all of these cytosolic DNA stimuli are of nuclear origin and released in the late stage of senescence. Here, we have demonstrated that cytosolic mtDNA release is essential in the early stage of senescence establishment across various senescent models, a previously unknown finding. Our study has elucidated the mechanisms governing mtDNA release, wherein nuclear translocation of EndoG, separated from AIF, allows the damaged mtDNA to bypass the mitochondrial membranes through VDACs. These mechanistic insights shed new light on how senescent cells-derived mtDNA engages senescence and inflammatory responses in cancer.

Furthermore, our investigation reveals that mtDNA released by senescent tumor cells enters the extracellular space, where it can enhance the immunosuppressive function of PMN-MDSCs. Remarkably, pharmacological inhibition of VDAC1 oligomerization using VBIT-4 significantly reduced the release of cytosolic mtDNA and SASP factors from senescent cells. This highlights the potential of this compound as a senostatic drug in cancer therapy. Furthermore, VBIT-4 treatment also resulted in a significant decrease in the uptake of extracellular mtDNA by PMN-MDSCs, consistent with the blockage of cytosolic mtDNA release. Decreased mtDNA release, in turn, enhanced the efficacy of chemotherapy by alleviating immunosuppression in the TME.

Collectively, our work provides novel insights into senescence, highlighting the dual role of accumulated senescent cell-associated mtDNA as an intracellular and extracellular signal molecule. Notably, our study establishes the clinical relevance of targeting mtDNA release during TIS, as evidenced by the correlation between elevated mtDNA levels and poor disease outcomes in patients with prostate cancer. These findings suggest that blocking mtDNA release in TIS holds great potential as a promising therapeutic strategy to enhance the efficacy of chemotherapy in clinical cancer treatment.

## Methods and materials

### Mice

All mice were maintained under specific pathogen-free conditions, and experiments were approved by the local ethical committee (‘‘Dipartimento della Sanità e Socialità, Esperimenti su animals,’’ Canton Ticino), authorization number 34293. *Mb21d1*^-/-^ (cGAS knockout) mice were a gift from Prof. Andrea Ablasser (Ecole Polytechnique Fédérale de Lausanne, Switzerland). For allograft experiments, C57BL6/N were challenged with 2.5×10^6^ TRAMP-C1 cells or RapidCap cells and then orally given Palbociclib (150 mg/Kg) three times and/or VBIT-4 (20 mg/Kg) five times per week when tumors were approximately 100 mm^3^. For the xenograft experiment, NRG mice were challenged with 2.5×10^6^ PC3 cells and then orally given Palbociclib when tumors were approximately 100 mm^3^. Tumor volume and survival rate were analyzed, and then sacrificed the mice when the tumor reached approximately 1000 mm^3^.

### Generation of *Pten* KO MEFs and *HRasV12* MEFs

Primary Mouse Embryonic Fibroblasts (MEF) were obtained from pregnant *Pten^flox/flox^* mice at 13.5 days post-coitum. Embryos were harvested, and the individual MEFs were cultured in DMEM containing 10% fetal bovine serum (FBS) and 1% Penicillin/Streptomycin (P/S). To prepare lentiviral particles, HEK-293T cells were transfected using JetPRIME® transfection reagents (JetPRIME^®^, Polyplus transfection, 114-07/712-60) as the manufacturer’s instructions. Primary MEFs were infected with retroviruses expressing either pWZL-Hygro (Addgene, #18750), pWZL-Hygro-CRE, or pWZL-Hygro-HRasV12 (Addgene, #18749) and selected with Hygromycin B (Invitrogen, 10687010) at a concentration of 50 μg/ml.

### Generation of *shEndog* and sh*Vdac1*, *2* MEFs

Primary MEFs were infected with shRNA using the mouse *Endog*-directed shRNA (Sigma), *Vdac1*, and *Vdac2* shRNA (Sigma). To prepare lentiviral particles, HEK-293T cells were transfected using JetPRIME transfection reagents as described above. MEFs cells were infected with the filtered lentiviral supernatant obtained from transfected HEK-293T cells and were subsequently selected using Puromycin (Sigma) at 3 μg/ml.

### Cell lines

PC3 cells were purchased from ATCC and cultured in RPMI 1640 supplemented with 10% FBS and 1% P/S. TRAMP-C1 and RM-1cells were purchased from ATCC; RapidCap is a kind gift from Prof. Lloyd C. Trotman (Cold Spring Harbor Laboratory) and cultured in DMEM supplemented with 10% FBS and 1% P/S. All cell lines were kept under controlled temperature (37°C) and CO_2_ (5%) and tested negative for Mycoplasma.

### Differentiation of BM-MDSCs in vitro

Mouse BM-MDSCs were differentiated in vitro as previously described^44^. In brief, bone marrow cells were flushed from the legs of C57BL/6 with RPMI 1640 medium. The cell pellets were lysed with ACK buffer (Gibco, A1049201) and resuspended in RPMI containing 40 ng/ml GM-CSF, 40 ng/ml IL-6, 10% FBS, and 1% P/S. On day 4, the cells were harvested and resuspended in the fresh RPMI 1640 containing 10% FBS and 1% P/S.

### In vitro T cell suppression assay

In vitro suppression assays were determined as previously described^44^. In brief, naïve splenocytes were labeled with 2.5 μM CFSE (Thermo Scientific, C34554) and activated in vitro with anti-CD3/CD28 beads (Gibco, 11452D) according to the manufacturer’s instructions. BM-MDSCs were added to the culture. After three days, the proliferation of CFSE-labelled CD8^+^ T cells and CD4^+^ T cells were analyzed by BD FACS Canto I.

### Senescence associated β-galactosidase (SA-β-gal) assay

For tissue-specific SA-β-gal assay, tumor samples were immediately frozen in OCT solution at −80 °C, and sections of 8 mm were prepared. Senescence-associated SA-β-gal staining was performed using Senescence β-Galactosidase Staining Kit (Cell Signaling Technology, 9860) according to the manufacturer’s instructions. Counterstaining was performed using Eosin staining (Alcohol-based Diapath, C0352). SA-β-gal staining was performed using Senescence β-Galactosidase Staining Kit (Cell Signalling Technology, Cat. No 9860) according to the manufacturer’s instructions.

### Immunohistochemistry (IHC) and Immunofluorescence (IF)

Tumor tissue samples were fixed in 10% neutral-buffered formalin (Thermo Scientific, Cat No. 5701) overnight and then washed thoroughly under running tap water, followed by processing using ethanol and embedded in paraffin according to standard protocols. Sections (5 mm) were prepared for antibody detection and hematoxylin and eosin staining. IHC tissue sections were processed as follows: de-paraffinized, unmasked, pre-staining, blockings, and secondary staining ^45^. Images were scanned with Aperio and opened with ImageScope v12.3.2.8013 (Leica Biosystem).

For in vitro immunofluorescence, all microscopy image cells were seeded on coverslips one day before fixation. After washing in PBS, cells were fixed with 4% paraformaldehyde for 20 min, permeabilized with 0.1% Triton X-100 in PBS for 15 min, blocked with PBS containing 10% FBS for 30 min, stained with primary antibodies at 4 ℃ for overnight, and stained with secondary antibodies for 60 min at room temperature. Cells were washed with 0.1% PBS-Tween 20 between each step. Slides were covered with mounting media with DAPI (Invitrogen, P36931), ready to be visualized under the fluorescent confocal microscope. Images were acquired on Leica TCS SP5 and merged using ImageJ software (NIH).

### Characterization of the immune tumor microenvironment

Tumors were isolated and digested in collagenase D and *DNase* I for 30 min at 37 °C to obtain a single-cell suspension. The single-cell were stained with Fixable viability stain (ThermoFisher Scientific) for 1 hour, and then CD16/ CD32 antibody was used to block the unspecific binding. Single-cell suspensions were stained with specific monoclonal antibodies (primary antibodies directly conjugated) to assess the phenotype and diluted 1:200. The antibodies used were: CD45 (clone 30-F11), Ly6C (clone HK1.4), Ly6G (clone 1A8), CD11b (clone M1/70), F4/80 (clone BM8), CD11c (clone N418), CD8 (clone 53-6.7), CD4 (clone GK1.5), CD3 (clone 17A2), B220 (clone RA3-6B2), CD19 (clone 1D3), CD49b (clone DX5). All antibodies were purchased from eBioscience, Biolegend, R&D, or BD. Samples were acquired on a BD FACSymphony (BD Biosciences). Data were analyzed using FlowJo software (TreeStar).

### Quantitative real-time PCR (RT-qPCR)

Total RNA was extracted with TRIzol reagent (Ambion, 15596026) following the manufacturer’s instructions. cDNA was obtained using an ImPROM II kit (Promega, A3800) according to the manufacturer’s instructions. RT-qPCR was performed using Gotaq® qPCR Master Mix, Promega® (A6002) on Step One Real-Time PCR systems (Applied Biosystems). The target genes were normalized to the housekeeping gene (β-Actin) shown as 2^−ΔΔCt^. The used primers are as follows: Cdkn1a (p21) forward 5’-TTCCCTCACAGGAGCAAAGT-3’, reverse 5-CGGCGCAACTGCTCACT-3’; Serpine1 (Pai1) forward 5’-TTCAGCCCTTGCTTGCCTC-3’, reverse 5’-ACACTTTTACTCCGAAGTCGGT-3’; 18s rDNA forward 5’-TAGAGGGACAAGTGGCGTTC-3’; 18s rDNA reverse5’-CGCTGAGCCAGTCAGTGT-3’; Tert forward 5’-CTAGCTCATGTGTCAAGACCCTCTT-3’, reverse 5’-GCCAGCACGTTTCTCTCGTT-3’; Dloop forward 5’-TCCTCCGTGAAACCAACAA-3’, reverse 5’-AGCGAGAAGAGGGGCATT-3’; IL8 forward 5’-CTGGTCCATGCTCCTGCTG-3’, reverse 5’-GGACGGACGAAGATGCCTAG-3’; Mmp3 forward 5’-TGGAGCTGATGCATAAGCCC-3’, reverse 5’-TGAAGCCACCAACATCAGGA-3’; Endog forward 5’-TTCCGCGAGGATGACTCTGT-3’, reverse 5’-CACCTGAGGCGCTACGTTG-3’; ACTB-forward 5’-GATGCACAGTAGGTCTAAGTGGAG-3’, reverse 5’-CACTCAGGGCAGGTGAAACT-3’; β-Actin forward 5’-GGCTGTATTCCCCTCCATCG-3’, reverse 5’-CCAGTTGGTAACAATGCCATG-3’; Ccl2 forward 5’-TTAAAAACCTGGATCGGAACCAA-3’, reverse 5’-GCATTAGCTTCAGATTTACGGGT-3’; Cxcl10 forward 5’-CCAAGTGCTGCCGTCATTTTC-3’, reverse 5’-GGCTCGCAGGGATGATTTCAA-3’; Ifi44 forward 5’-CTGATTACAAAAGAAGACATGACAGAC-3’, reverse 5’-AGGCAAAACCAAAGACTCCA-3’; Ifit1 forward 5’-CAAGGCAGGTTTCTGAGGAG-3’, reverse 5’-GACCTGGTCACCATCAGCAT-3’; Ifnb forward 5’-CCCTATGGAGATGACGGAGA-3’, reverse 5’-CCCAGTGCTGGAGAAATTGT-3’; Isg15 forward 5’-CTAGAGCTAGAGCCTGCAG-3’, reverse 5’-AGTTAGTCACGGACACCAG-3’; Vdac1 forward 5’-ACTAATGTGAATGACGGGACA-3’, reverse 5’-GCATTGACGTTCTTGCCAT-3’; Cox1 forward 5’-GCCCCAGATATAGCATTCCC-3’, reverse 5’-GTTCATCCTGTTCCTGCTCC-3’; IL1b forward 5’-TGTAATGAAAGACGGCACACC-3’, reverse 5’-TCTTCTTTGGGTATTGCTTGG-3’; Tnfa forward 5’-CCCTCACACTCAGATCATCTTCT-3’, reverse 5’-GCTACGACGTGGGCTACAG-3’; IL12 forward 5’-TACTAGAGAGACTTCTTCCACAACAAGAG-3’, reverse 5’-TCTGGTACATCTTCAAGTCCTCATAGA-3’;S100A8 forward 5’-GTCCTCAGTTTGTGCAGAATATAAA-3’, reverse 5’-GCCAGAAGCTCTGCTACTCC-3’; IL6 forward 5’-TAGTCCTTCCTACCCCAATTTCC-3’, reverse 5’-TTGGTCCTTAGCCACTCCTTC-3’; Nos2 forward 5’-GTTCTCAGCCCAACAATACAAGA-3’, reverse 5’-GTGGACGGGTCGATGTCAC-3’; Arg1 forward 5’-CCACACTGACTCTTCCATTCTT-3’, reverse 5’-GATTATCGGAGCGCCTTTCT-3’; hLINE1-forward 5’-TCACTCAAAGCCGCTCAACTAC-3’, reverse 5’-TCTGCCTTCATTTCGTTATGTACC-3’; hNd1 forward 5’-ATACCCATGGCCAACCTCCT-3’, reverse 5’-GGGCCTTTGCGTAGTTGTAT-3’.

### Cytosolic mitochondrial DNA (mtDNA) extraction and quantification

1×10^6^ Cells were each divided into two equal aliquots, and one aliquot was resuspended in 300 μl of 50 μM NaOH and boiled at 95℃ for 30 min to solubilize DNA. 30 μl of 1 M Tris-HCl pH 8 was added to neutralize the pH, and these extracts served as normalization controls for total mtDNA. The second equal aliquots were resuspended in roughly 300 μl buffer containing 150 mM NaCl, 50 mM HEPES pH 7.4, and 25 μg/ml digitonin (EMD Chemicals). The homogenates were incubated on a rotator for 10 min at room temperature, followed by centrifugation at 980g 4℃ for 3 min three times to pellet intact cells. The cytosolic supernatants were then spun at 17000g for 20 min to pellet any remaining cellular debris. Quantitative PCR was performed on both whole-cell extracts and cytosolic fractions using nuclear DNA primers (Tert) and mtDNA primers (Dloop3, Cox1), and the CT values obtained for mtDNA abundance for whole-cell extracts served as normalization controls for the mtDNA values obtained from the cytosolic fractions.

### Western blot

Cells were lysed using 1×RIPA buffer (Cell signaling, 9806) containing protease and phosphatase inhibitors (Thermo Fisher, A32959) and then incubated on ice for 30 min. Samples were centrifuged at 13,000 g for 10 min. Protein concentration was determined by the BCA kit (Thermo Fisher, 23227). Equal amounts of proteins were subjected to SDS-polyacrylamide gel electrophoresis (SDS-PAGE), 10%, and transferred onto 0.45 µm PVDF membrane (Thermo Scientific, 88518). Membranes were blocked in 5% milk solution after protein transfer. Membranes were probed with the indicated antibodies overnight at 4 °C. The primary antibodies are: p16 (Abcam, ab211542), p21 (Abcam, ab107099), p27 (CST, 3698S), HSP90 (CST, 4877S), PTEN (CST, 9188S), RAS (BD Transduction, 610001), p-HP1γ (CST, 2600S), p-KAP1 (Thermo Fisher, A300-767A), p-γH2A.X (CST, 9718S), GAPDH (CST, 5174S), H4 (CST, 2592S), p-STING (CST, 72971S), STING (CST, 13647), p-TBK1 (CST, 5483S), TBK1 (CST, 3504), STAT1 (CST, 9172), p-STAT1 (Y701) (CST, 9167), cGAS (CST, 31659), ENDOG (Sigma, SAB3500213), AIF (CST, 5318T), VDAC1 (CST, 4661), VDAC2 (CST, 9412S). After washing with 1×PBST, the membranes were incubated with horseradish peroxidase-conjugated (HRP-linked) secondary antibodies anti-rat IgG (ThermoFisher Scientific, 31470, 1:5000), anti-rabbit IgG (Promega, W4011, 1:5000) or anti-mouse IgG (Promega, W4021, 1:5000) and developed using enhanced chemiluminescence (ECL) substrate (Thermo Scientific, 32106). Membranes were exposed to the FusionSolo S imaging system (Vilber). Blots were semi-quantitatively analyzed by densitometry using ImageJ 1.52 v (National Institutes of Health).

### Cell fractionation

A method was carried out for nuclei isolation as previously described with some modifications (Gagnon et al., 2014). Briefly, harvested cells were washed in ice-cold PBS and resuspended in hypotonic lysis buffer (10 mM Tris, pH 7.5, 10 mM NaCl, 3 mM MgCl_2_, 0.3% NP-40, and 10% glycerol) in the presence of a protease/phosphatase inhibitor cocktail (Thermo Fisher Scientific). The suspension was incubated for 30 min on ice, passed ten times through a 28-gauge blunt-ended needle, and centrifuged at 400 g for 5 min at 4 ℃. The nuclear pellet was washed three times with hypotonic lysis buffer and collected as a total nuclear fraction.

To isolate mitochondria-enriched cellular fractions, a crude mitochondrial fraction was first obtained, as described previously, with minor modifications (Frezza et al., 2007). In brief, cells were washed with ice-cold PBS and suspended in chilled mitochondria isolation buffer (IBc) (10 mM Tris-MOPS, pH 7.4, 10 mM EGTA-Tris, 200 mM sucrose, and 5 mM MgCl_2_) with a protease/phosphatase inhibitor cocktail. The cells were homogenized in IBc buffer using a Teflon pestle (about 50 strokes) and centrifuged at 500 g for 10 min at 4℃, followed by further centrifuging at 2,000 g for 10 min at 4℃ to remove unbroken cells, cell debris, and nucleus. Then the supernatant was collected and centrifuged at 7,500 g for 10 min at 4℃. The pellets containing mitochondria were washed twice with IBc buffer and saved the pellets as mitochondria fraction. Mitochondria-free cytosolic fraction was collected from the supernatant and centrifuged at 16,000 g for 15 min at 4℃. The supernatant was saved as a cytosolic fraction. Protein levels were determined using a Pierce BCA protein assay (Thermo Fisher Scientific), and each fraction was loaded for western blotting to confirm purity.

### cGAS Immunoprecipitation-PCR

To pull down cytosolic endogenous cGAS, MEFs were fixed for 10 min in 4% PFA and quenched with 1 M Tris pH 7.4 for 5 min. Cytosolic fractions were isolated from the fixed cells following the previous procedures. Cytoplasm lysates were precleared with protein G magnetic beads (Invitrogen) and immunoprecipitated using anti-Rb cGAS (CST) or Rb Ig (CST) Protein G beads at 4℃ overnight. Immunoprecipitates were washed five times with 50 mM Tris, 150 mM NaCl, 1 mM EDTA, and 0.05% NP-40 (Wash Buffer). Eluted DNA was reverse cross-linked and treated with 0.2 mg/ml proteinase K (Qiagen) for 2 h at 60℃ and heat inactive at 95 for 15 min. DNA was extracted with QIAquick PCR purification kit (Qiagen). DNA elution was used in qPCR analysis to measure the abundance of specific DNA sequences.

### Co-culture Experiment

PC3 cells were treated with vehicle, Palbociclib (10 μM), and Docetaxel (10 nM) for 2 days. Then, cells were washed and re-seeded in 24-wells plates at 50,000 seeding density. 2×10^5^ BM-MDSC were seeded into a 0.4 µm cell culture insert (Falcon, 353495) and co-cultured with PC3 cells in the bottom chamber. Three days post-co-culturing, BM-MDSC cells were collected, and human mtDNA was analyzed by quantitative PCR with human mtDNA primers (ND1) and normalized to mouse nuclear DNA (Tert).

### DNA dot blot

DNA samples were denatured at 99°C for 10 min and chilled on ice for 5 min and then spotted onto Hybond-N1 nitrocellulose membranes (ThermoFisher) under vacuum using a 96-well Dot Blot hybridization manifold (BioRad, BIO-DOT Apparatus). The membrane was washed twice in 2× SSC buffer and dried for 1 h at 80°C. After ultraviolet cross-linking, membranes were blocked with 10% non-fat milk and 1% BSA in PBT (PBS 1 0.1% Tween20) for 1h, followed by 5mC antibody (Abcam, ab10805) (1:1000) incubation overnight at 4°C. Membranes were washed four times with PBST and incubated for 1h with HRP-conjugated anti-rabbit secondary antibody. Following treatment with enhanced chemiluminescence substrate, membranes were scanned on the FusionSolo S imaging system (Vilber). To control for loading, membranes were stained with 0.02% methylene blue solution.

### Extraction and quantification of plasma cfDNA

cfDNA was extracted from 600 µL of conditioned medium and 100 µL of human/murine plasma using MagMAX Cell-Free DNA Isolation Kit (ThermoFisher Scientific, A29319). The concentration of cfDNA was measured using Qubit dsDNA HS Assay Kit with Qubit 4.0 Fluorometer (ThermoFisher Scientific). The absolute copy numbers were measured using quantitative PCR using indicated primers. The copy number was calculated using the standard curves for cloned Tert and Dloop in a pMD19-T Easy vector (TaKaRa, 3271), respectively. Primer sequences are shown above.

### Gene Expression Profiling

RNA sequencing was performed at the Institute of Oncology Research using the NEBNext Ultra Directional II RNA library preparation kit for Illumina and sequenced on the Illumina NextSeq500 with single-end, 75 base pair long reads. The overall quality of sequencing reads was evaluated using FastQC. STAR (v.2.7.10a)^46^ was used to sequence alignments to the reference mouse genome (GRCm39). Gene Transfer File (GTF) vM27 by Gencode was used to quantify gene expression at gene level. Further analysis were performed in R Statistical environment (v.4.1.0). Genes without counts were removed for the analysis and differential expression analysis was performed using DESeq2. In DESeq2 function the parameter independent Filtering was set up to TRUE to remove genes with low mean normalized counts. Pathway analysis was performed using Camera^47^ and custom gene signature of MDSC functions from different works 22348096, 32086381, 33526920, 27381735, 31533831. All graphical representations were edited using ggplot2 and pheatmap functions.

### Statistical analysis

All values are expressed as the mean and SEM. Statistical analysis was performed with the unpaired t-test for two groups or one-way ANOVA (GraphPad Software) used for multiple groups, with all data points showing a normal distribution. A two-way ANOVA was used for experiments with two independent variables, in combination. The researchers were not blinded to the allocation of treatment groups when performing experiments and data assessment. Sample sizes were selected based on preliminary results to ensure adequate power. *p* values < 0.05 were considered significant.

### Data availability statement

All datasets have been deposited in the Genome Sequence Archive in the National Genomics Data Center with accession numbers as indicated.

## Acknowledgments

We acknowledge Prof. A. Alimonti’s laboratory members for the scientific discussions. We also thank Dr. David Jarrossay of the Bios^+^ FACS facility and all the genomics and animal core facility members for technical assistance. We acknowledge all the patients that participated in the study protocols. We also thank Prof. Andrea Ablasser for kindly providing *Mb21d1*^-/-^ (cGAS knockout) mice and Prof. Georg Häcker for sharing pF5UAS_UL12.5_Puro and pFU-GEV16 plasmids. This work was supported by the Horten Foundation, Dr. Josef Steiner Foundation (Dr. Josef Steiner Cancer Research Prize 2015), Swiss Cancer Research Foundation (KFS-4267-08-2017, KFS-5262-02-2021, and RWP-4813-06-2019), SNSF (grants 310030_176045, 310030B_201274, and CRSII5_202302), Novartis Foundation, ISREC Foundation, AIRC (IG 22030), PCF (19CHAL08), and ERC (CoG 683136).

## Contributions

P.L., L.L., J.C., and A.A. conceived the ideas. P.L., L.L. and A.A. designed the experiments and wrote the manuscript. L.P. performed most of the experiments and analyzed the data. The flow cytometry assays data, and IHC quantification was analyzed by L.L. and N.B.. RNA sequencing was performed by A.R., and M.T. performed the bioinformatics analysis. G.A. and E.M. took care of transgenic mouse model husbandry. S.M. performed immunohistochemistry. B.C. and P.L. performed the immunofluorescence staining and data analysis. M.D. performed the cfDNA isolation and mtDNA quantification in patients’ plasma samples. All authors contributed to the revision of the manuscript.

## Materials & Correspondence

Please address all correspondence and requests to Professor Andrea Alimonti.

## Reference

1. Calcinotto, A., et al. IL-23 secreted by myeloid cells drives castration-resistant prostate cancer. Nature (2018).

2. Demaria, M., et al. Cellular Senescence Promotes Adverse Effects of Chemotherapy and Cancer Relapse. Cancer Discovery 7, 165–176 (2017).

3. Dou, Z., et al. Cytoplasmic chromatin triggers inflammation in senescence and cancer. Nature 550, 402–406 (2017).

4. De Cecco, M., et al. L1 drives IFN in senescent cells and promotes age-associated inflammation. Nature 566, 73–78 (2019).

5. Liu, X., et al. Resurrection of endogenous retroviruses during aging reinforces senescence. Cell 186, 287–304 e226 (2023).

6. Yum, S., Li, M., Fang, Y. & Chen, Z.J. TBK1 recruitment to STING activates both IRF3 and NF-kappaB that mediate immune defense against tumors and viral infections. Proc Natl Acad Sci U S A 118(2021).

7. McArthur, K., et al. BAK/BAX macropores facilitate mitochondrial herniation and mtDNA efflux during apoptosis. Science 359(2018).

8. Riley, J.S., et al. Mitochondrial inner membrane permeabilisation enables mtDNA release during apoptosis. EMBO J 37(2018).

9. Sun, B., et al. Dengue virus activates cGAS through the release of mitochondrial DNA. Sci Rep 7, 3594 (2017).

10. Ni, G., Ma, Z. & Damania, B. cGAS and STING: At the intersection of DNA and RNA virus-sensing networks. PLoS Pathog 14, e1007148 (2018).

11. West, A.P., et al. Mitochondrial DNA stress primes the antiviral innate immune response. Nature 520, 553–557 (2015).

12. Shimada, K., et al. Oxidized mitochondrial DNA activates the NLRP3 inflammasome during apoptosis. Immunity 36, 401–414 (2012).

13. Miller, K.N., et al. Cytoplasmic DNA: sources, sensing, and role in aging and disease. Cell 184, 5506–5526 (2021).

14. White, M.J., et al. Apoptotic caspases suppress mtDNA-induced STING-mediated type I IFN production. Cell 159, 1549–1562 (2014).

15. Hajizadeh, S., DeGroot, J., TeKoppele, J.M., Tarkowski, A. & Collins, L.V. Extracellular mitochondrial DNA and oxidatively damaged DNA in synovial fluid of patients with rheumatoid arthritis. Arthritis Res Ther 5, R234–240 (2003).

16. Ryu, C., et al. Extracellular Mitochondrial DNA Is Generated by Fibroblasts and Predicts Death in Idiopathic Pulmonary Fibrosis. Am J Respir Crit Care Med 196, 1571–1581 (2017).

17. Iske, J., et al. Senolytics prevent mt-DNA-induced inflammation and promote the survival of aged organs following transplantation. Nat Commun 11, 4289 (2020).

18. Palmbos, P.L., et al. Clinical outcomes and markers of treatment response in a randomized phase II study of androgen deprivation therapy with or without palbociclib in RB-intact metastatic hormone-sensitive prostate cancer (mHSPC). J Clin Oncol 38(2020).

19. Palmbos, P.L., et al. A Randomized Phase II Study of Androgen Deprivation Therapy with or without Palbociclib in RB-positive Metastatic Hormone-Sensitive Prostate Cancer. Clin Cancer Res 27, 3017–3027 (2021).

20. Moiseeva, O., Bourdeau, V., Roux, A., Deschenes-Simard, X. & Ferbeyre, G. Mitochondrial dysfunction contributes to oncogene-induced senescence. Mol Cell Biol 29, 4495–4507 (2009).

21. Spadafora, D., Kozhukhar, N., Chouljenko, V.N., Kousoulas, K.G. & Alexeyev, M.F. Methods for Efficient Elimination of Mitochondrial DNA from Cultured Cells. PLoS One 11, e0154684 (2016).

22. Gluck, S., et al. Innate immune sensing of cytosolic chromatin fragments through cGAS promotes senescence. Nat Cell Biol 19, 1061–1070 (2017).

23. Huang, K.J., Ku, C.C. & Lehman, I.R. Endonuclease G: a role for the enzyme in recombination and cellular proliferation. Proc Natl Acad Sci U S A 103, 8995–9000 (2006).

24. Wiehe, R.S., et al. Endonuclease G promotes mitochondrial genome cleavage and replication. Oncotarget 9, 18309–18326 (2018).

25. Kim, J., et al. VDAC oligomers form mitochondrial pores to release mtDNA fragments and promote lupus-like disease. Science 366, 1531–1536 (2019).

26. Correia-Melo, C. & Passos, J.F. Mitochondria: Are they causal players in cellular senescence? Biochim Biophys Acta 1847, 1373–1379 (2015).

27. Passos, J.F., et al. Mitochondrial dysfunction accounts for the stochastic heterogeneity in telomere-dependent senescence. PLoS Biol 5, e110 (2007).

28. Kaplon, J., et al. A key role for mitochondrial gatekeeper pyruvate dehydrogenase in oncogene-induced senescence. Nature 498, 109–112 (2013).

29. Korolchuk, V.I., Miwa, S., Carroll, B. & von Zglinicki, T. Mitochondria in Cell Senescence: Is Mitophagy the Weakest Link? EBioMedicine 21, 7–13 (2017).

30. Lin, J.L.J., et al. Oxidative Stress Impairs Cell Death by Repressing the Nuclease Activity of Mitochondrial Endonuclease G. Cell Rep 16, 279–287 (2016).

31. Ishihara, Y. & Shimamoto, N. Involvement of endonuclease G in nucleosomal DNA fragmentation under sustained endogenous oxidative stress. J Biol Chem 281, 6726–6733 (2006).

32. van Loo, G., et al. Endonuclease G: a mitochondrial protein released in apoptosis and involved in caspase-independent DNA degradation. Cell Death Differ 8, 1136–1142 (2001).

33. Wang, X., Yang, C., Chai, J., Shi, Y. & Xue, D. Mechanisms of AIF-mediated apoptotic DNA degradation in Caenorhabditis elegans. Science 298, 1587–1592 (2002).

34. Parrish, J., et al. Mitochondrial endonuclease G is important for apoptosis in C-elegans. Nature 412, 90–94 (2001).

35. Marcotte, R., Lacelle, C. & Wang, E. Senescent fibroblasts resist apoptosis by downregulating caspase-3. Mech Ageing Dev 125, 777–783 (2004).

36. Wang, L., Lankhorst, L. & Bernards, R. Exploiting senescence for the treatment of cancer. Nat Rev Cancer 22, 340–355 (2022).

37. Krtolica, A., Parrinello, S., Lockett, S., Desprez, P.Y. & Campisi, J. Senescent fibroblasts promote epithelial cell growth and tumorigenesis: a link between cancer and aging. Proc Natl Acad Sci U S A 98, 12072–12077 (2001).

38. Coppe, J.P., Desprez, P.Y., Krtolica, A. & Campisi, J. The senescence-associated secretory phenotype: the dark side of tumor suppression. Annu Rev Pathol 5, 99–118 (2010).

39. Di Mitri, D., et al. Tumour-infiltrating Gr-1+ myeloid cells antagonize senescence in cancer. Nature 515, 134–137 (2014).

40. Wang, G., et al. Targeting YAP-Dependent MDSC Infiltration Impairs Tumor Progression. Cancer Discov 6, 80–95 (2016).

41. Weber, R., et al. IL-6 as a major regulator of MDSC activity and possible target for cancer immunotherapy. Cell Immunol 359, 104254 (2021).

42. Veglia, F., Sanseviero, E. & Gabrilovich, D.I. Myeloid-derived suppressor cells in the era of increasing myeloid cell diversity. Nat Rev Immunol 21, 485–498 (2021).

43. Gorbunova, V., et al. The role of retrotransposable elements in ageing and age-associated diseases. Nature 596, 43–53 (2021).

44. Solito, S., et al. Methods to Measure MDSC Immune Suppressive Activity In Vitro and In Vivo. Curr Protoc Immunol 124, e61 (2019).

45. Guccini, I., et al. Senescence Reprogramming by TIMP1 Deficiency Promotes Prostate Cancer Metastasis. Cancer Cell 39, 68–82 e69 (2021).

46. Dobin, A., et al. STAR: ultrafast universal RNA-seq aligner. Bioinformatics 29, 15–21 (2013).

47. Wu, D. & Smyth, G.K. Camera: a competitive gene set test accounting for intergene correlation. Nucleic Acids Res 40, e133 (2012).

