## Supplemental figures for "Mitochondrial DNA released by senescent cells triggers immunosuppression in cancer"

### Extended Data Fig. 1

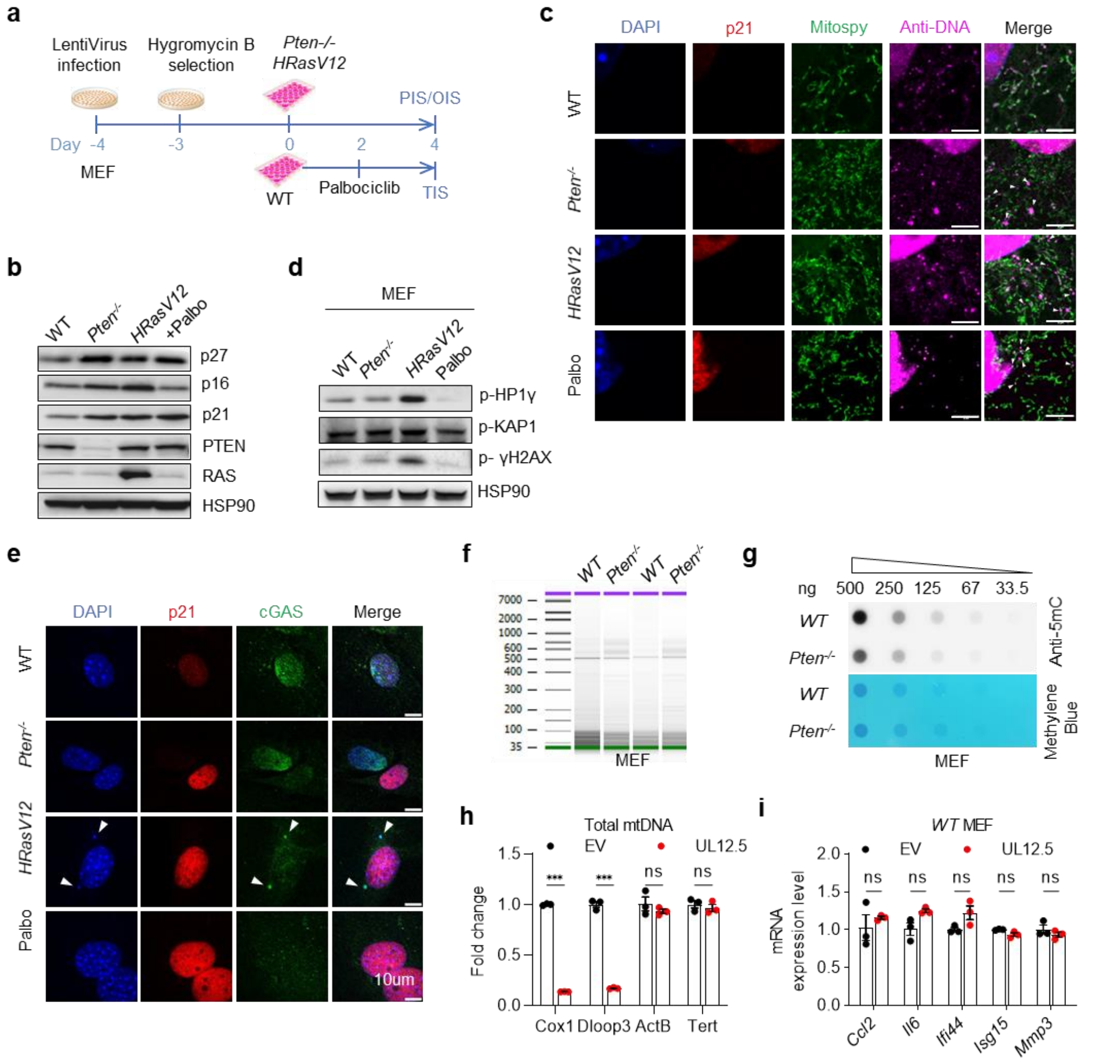


**Extended Data Fig. 1 Senescent cells accumulate fragmented and hypomethylated cytosolic mtDNA**

**a**, Scheme of timeline and experimental design of PIS, OIS, and TIS construction in *Pten*$^{fl/fl}$ MEFs. **b**, Western blot of senescence markers in senescent MEFs as indicated. **c**, Confocal microscopy images of indicated staining in senescent MEFs. White arrows indicate cytosolic DNA. Scale bar: 5 µm. **d**, Western blot analysis of DNA damage markers in senescent MEFs as indicated. **e**, Confocal microscopy images of indicated staining in senescent MEFs. White arrows indicate CCF in OIS. Scale bar: 10 µm. **f**, The size distribution of CM-DNA from *WT* and *Pten*$^{-/-}$ MEFs was analyzed by Bioanalyzer 2100. **g**, DNA Dot blot analysis using 5mC specific antibody to detect 5mC level on two-fold dilution of mtDNA from *WT* and *Pten*$^{-/-}$ MEFs. Methylene blue staining was used as a loading control. **h**, qPCR analysis of total mtDNA and nDNA in MEF cell after HSV1 UL12.5 or empty vector (EV) lentivirus infection. **i**, SASP genes expression was examined in *WT* MEFs were transduced with HSV1 UL12.5 or empty vector (EV). All values are presented as the mean ± SEM. Multiple unpaired t-test was used in **h** and **i**. *p < 0.05; **p < 0.01; ***p < 0.001; ns, not significant.

### Extended Data Fig. 2

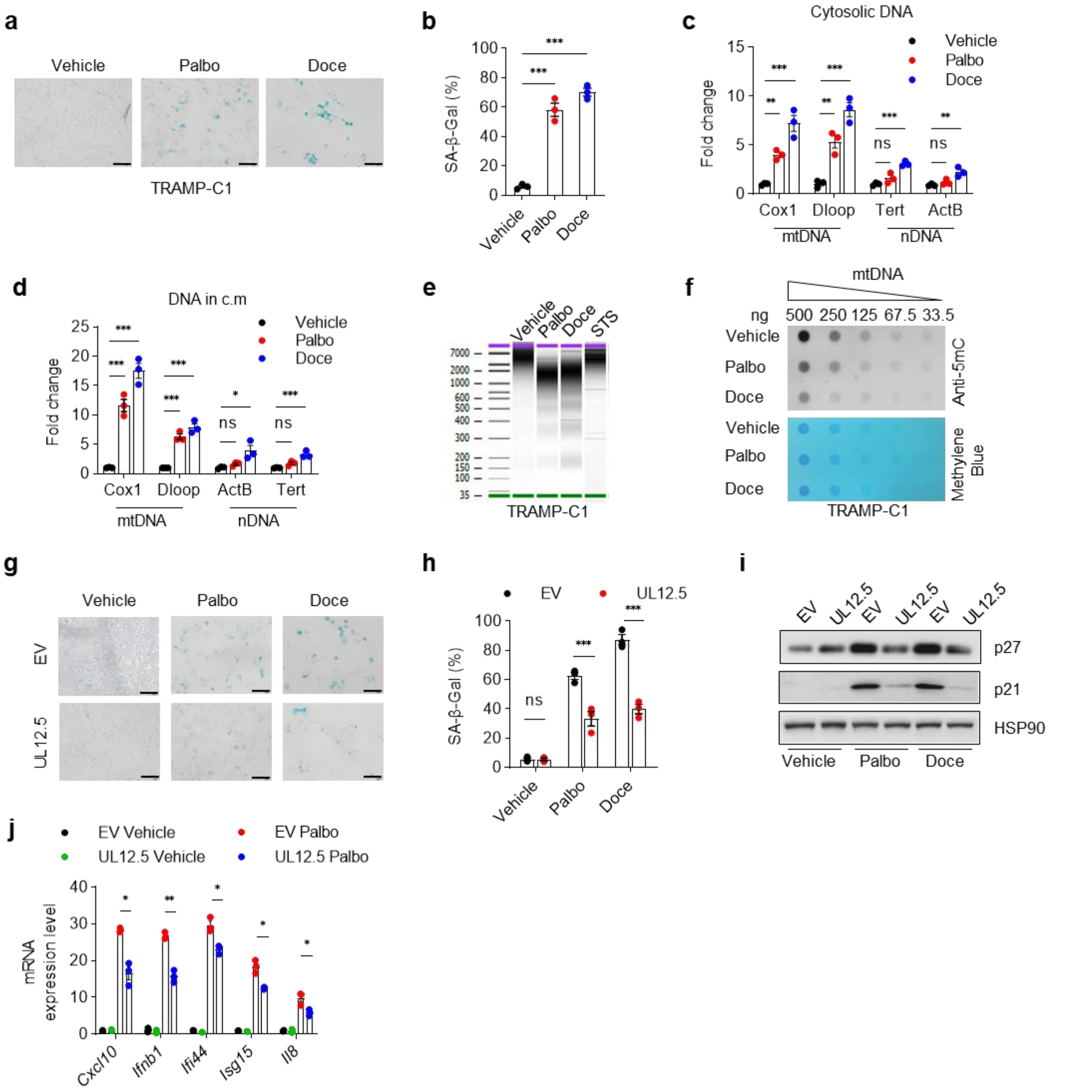


**Extended Data Fig. 2 Senescent tumor cells release mtDNA in the cytosol and extracellular space**

**a**, **b**, Representative images (**a**) and quantification (**b**) of SA-β-gal staining in TRAMP-C1 cell treated with 2.5 µM Palbociclib (Palbo) or 10 nM Docetaxel (Doce) for 48 h. Scale bar: 100 µm **c**, Quantification of cytosolic mtDNA and nDNA in TRAMP-C1 treated with Palbo and Doce. **d**, Quantification of mtDNA and nDNA in conditioned medium collected from TRAMP-C1 treated with Palbo and Doce as mentioned before. **e**, The size distribution of c.m-DNA from Palbo, Doce, or Staurosporine (STS) treated TRAMP-C1 cells were analyzed by Bioanalyzer 2100. **f**, DNA Dot blot analysis using anti-5mC antibody to detect 5mC level on two-fold dilution of mtDNA from Palbo or Doce treated TRAMP-C1 cells. Methylene blue staining was used as a loading control.

**g-j**, TRAMP-C1 cell were infected with HSV1 UL12.5 or empty vector (EV) lentivirus. SA-β-gal staining and quantification (**g**, **h**), protein (**i**), and SASP genes expression (**j**) were examined in TRAMP-C1 EV and TRAMP-C1 UL12.5 cells treated with Palbo or Doce. All values are presented as the mean ± SEM. Multiple unpaired t-test was used in **h**. One-way ANOVA followed by Tukey's multiple comparisons test was used in **b-d**. Two-way ANOVA followed by Dunnett's multiple comparisons test was used to evaluate the statistical significance in **j**. $*p < 0.05$; $**p < 0.01$; $***p < 0.001$; ns, not significant.

**Extended Data Fig. 3**

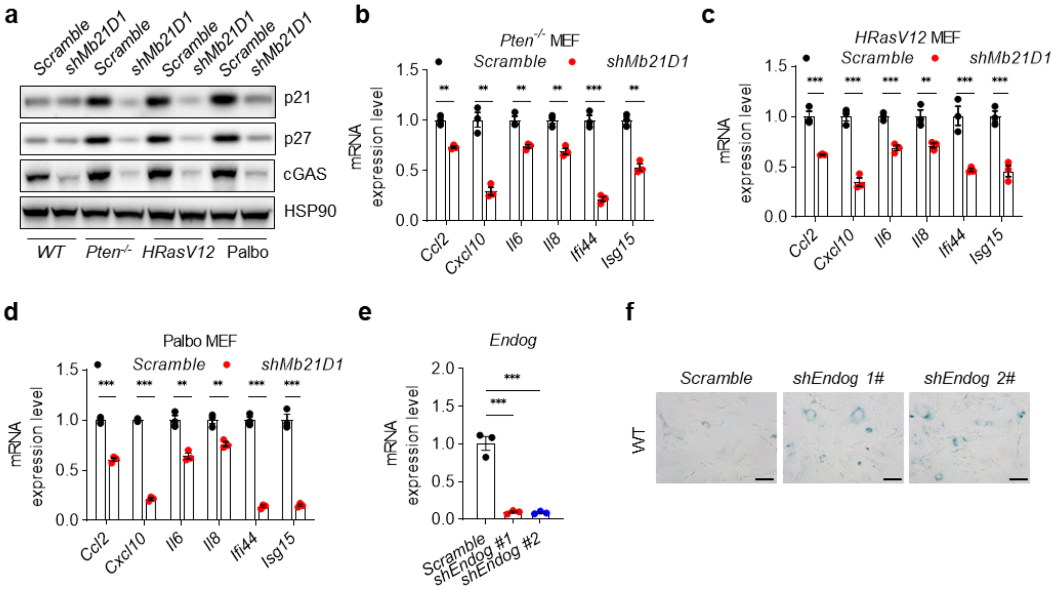


**Extended Data Fig. 3 Senescent cells release mtDNA into the cytoplasm and active cGAS-STING pathway**

**a**, MEFs were transduced with cGAS shRNA or scramble vectors, and senescence was induced as aforementioned. Immunoblot analysis of cGAS expression was shown. **b-d**, SASP gene expression was examined in the indicated models. **e**, RT-qPCR analysis of *Endog* mRNA expression level in MEFs after *shEndog* lentivirus infection. **f**, Representative images of SA-β-gal staining in *WT* and *Pten*$^{-/-}$ MEFs after *Endog* knocking down. All values are presented as the mean ± SEM. Multiple unpaired t-test was used in **b-d**. One-way ANOVA followed by Tukey's multiple comparisons test was used in **e**. *p < 0.05; **p < 0.01; ***p < 0.001; ns, not significant.

### Extended Data Fig. 4

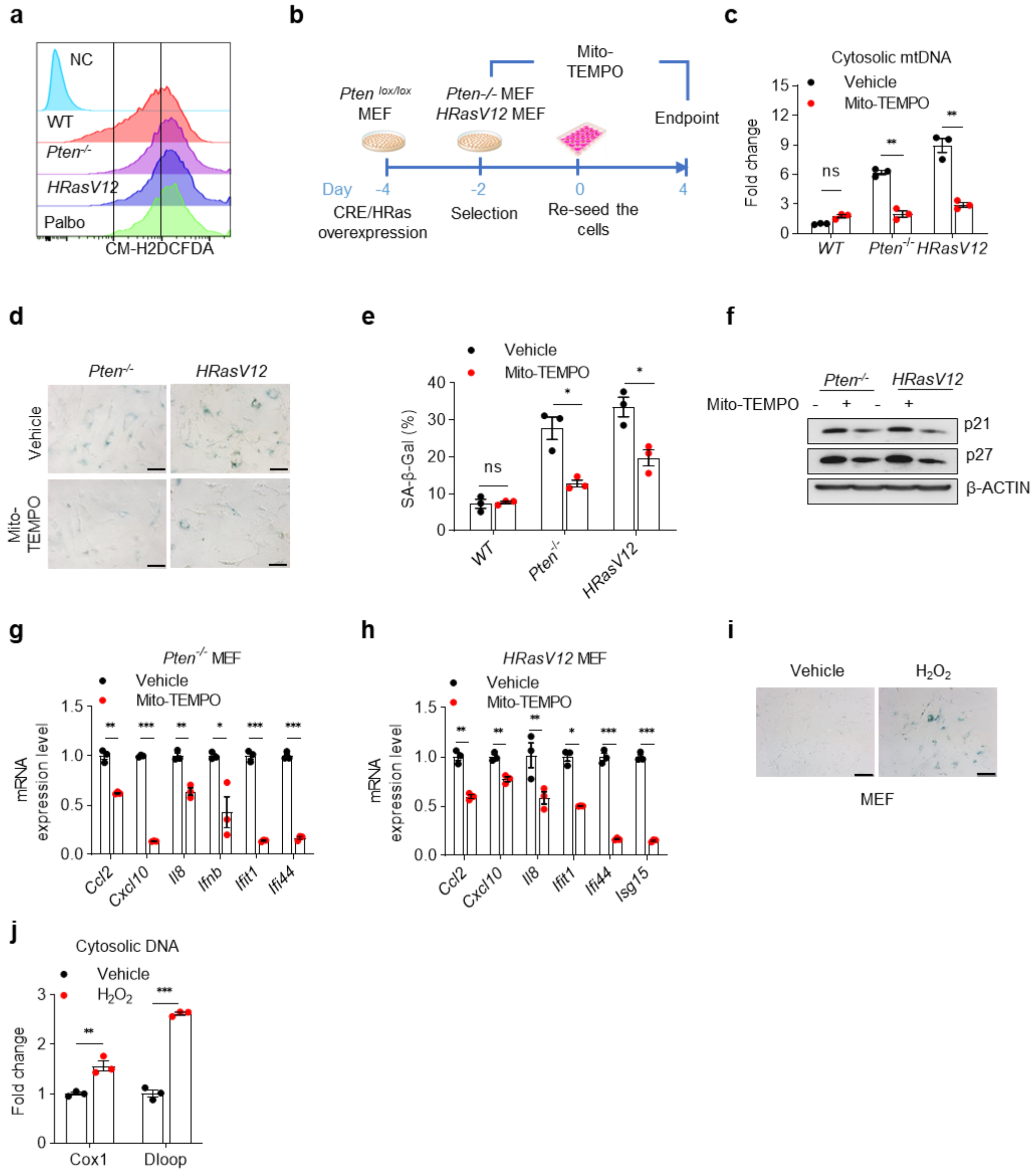


**Extended Data Fig. 4 mtROS induce mtDNA release in senescent cells**

**a**, Fluorescence intensity of senescent MEFs analyzed by flow cytometry after staining with CM-H2DCFDA. **b**, Timeline for Mito-TEMPO treatment in *Pten*$^{-/-}$ and *HRasV12* MEFs. **c**, Quantification of cytosolic mtDNA in senescent MEFs treated with 5 μM Mito-TEMPO. **d-h**, SA-β-gal staining (**d**), and quantification (**e**) indicated protein (**f**) and SASP genes expression (**g**, **h**) were detected in senescent MEFs treated with Mito-TEMPO. **i**, Representative images of SA-β-gal staining in *WT* MEFs treated with 40 μM $H_2O_2$ for 4 h and 16 h recovery. **j**, Quantification of cytosolic mtDNA in MEFs treated with $H_2O_2$ as aforementioned. All values are presented as the mean ± SEM.

Multiple unpaired t-test was used in **c**, **e**, **g, h** and **j**. $^{*}p < 0.05$; $^{**}p < 0.01$; $^{***}p < 0.001$; ns, not significant.

### Extended Data Fig. 5

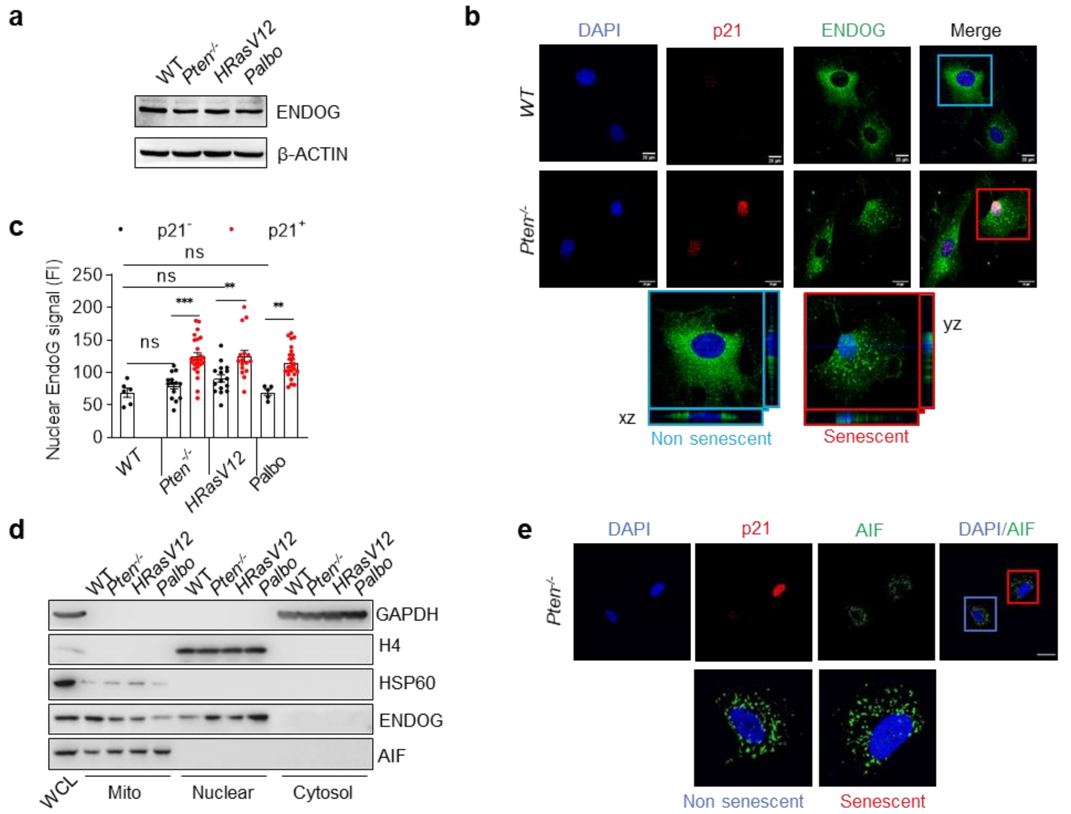


**Extended Data Fig. 5 Nuclear translocation of EndoG is required for mtDNA release**

**a**, Immunoblot analysis of EndoG in senescent MEFs. **b**, Confocal microscopy images of EndoG nuclear translocation in senescent MEFs. Scale bar: 20 μm. **c** Fluorescence intensity of nuclear-localized EndoG in senescent MEFs. **d**, WB analysis of EndoG and AIF cellular localization in senescent MEFs. WCL, whole cell lysate. **e**, Confocal microscopy images of AIF in *Pten*$^{-/-}$ MEFs. Scale bar: 20 μm. All values are presented as the mean ± SEM. Multiple unpaired t-test was used in **c**. $^{*}p < 0.05$; $^{**}p < 0.01$; $^{***}p < 0.001$; ns, not significant.

### Extended Data Fig. 6

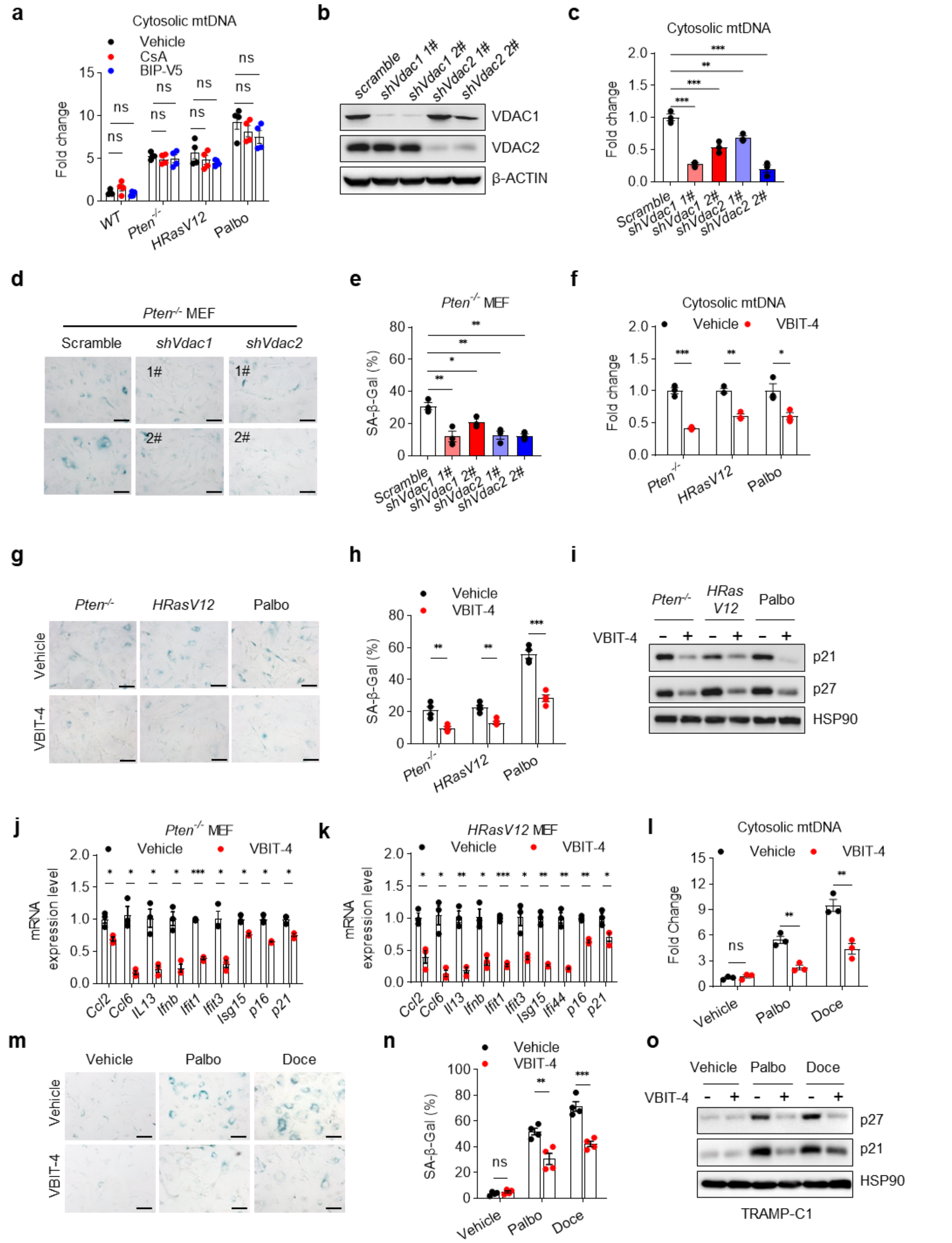


**Extended Data Fig. 6 Cytosolic mtDNA releases in senescent cells occurs through VDAC**

**a**, Quantification of cytosolic mtDNA in senescent MEFs treated with mPTP inhibitor CsA at 2.5 μM and BAX inhibitor BIP-V5 at 10 μM. **b,** WB analysis of VDAC1 and VDAC2 in MEFs infected with *Scramble* or *shVdac1*, *shVdac2* lentivirus. **c**,

Quantification of cytosolic mtDNA in *Pten*$^{-/-}$ MEF infected *Scramble* or *shVdac1*, *shVdac2* lentivirus. **d-e**, Representative images (**d**) and quantification (**e**) of SA-β-gal staining of *Pten*$^{-/-}$ MEF infected *Scramble* or *shVdac1*, *shVdac2* lentivirus. **f**, Quantification of cytosolic mtDNA in senescent MEFs treated with VDAC1 inhibitor VBIT-4 at 5 μM. **g-i**, SA-β-gal staining (**g**), and quantification (**h**), indicated protein (**i**) was detected in senescent MEFs treated with VBIT-4. **j**, **k** Indicated SASP and senescence marker genes expression in *Pten*$^{-/-}$ (**j**) and *HRasV12* (**k**) MEFs treated with VBIT-4. **l**, Quantification of cytosolic mtDNA in senescent TRAMP-C1 cells treated with VBIT-4 at 5 μM. **m-n**, SA-β-gal staining (**m**), and quantification (**n**) indicated protein (**o**) were detected in senescent TRAMP-C1 treated with VBIT-4. All values are presented as the mean ± SEM. Multiple unpaired t-test was used in **f**, **h**, **j-l**, **n**. One-way ANOVA followed by Tukey's multiple comparisons test was used in **a**, **c**, **e**. $*p < 0.05$; $**p < 0.01$; $***p < 0.001$; ns, not significant.

### Extended Data Fig. 7

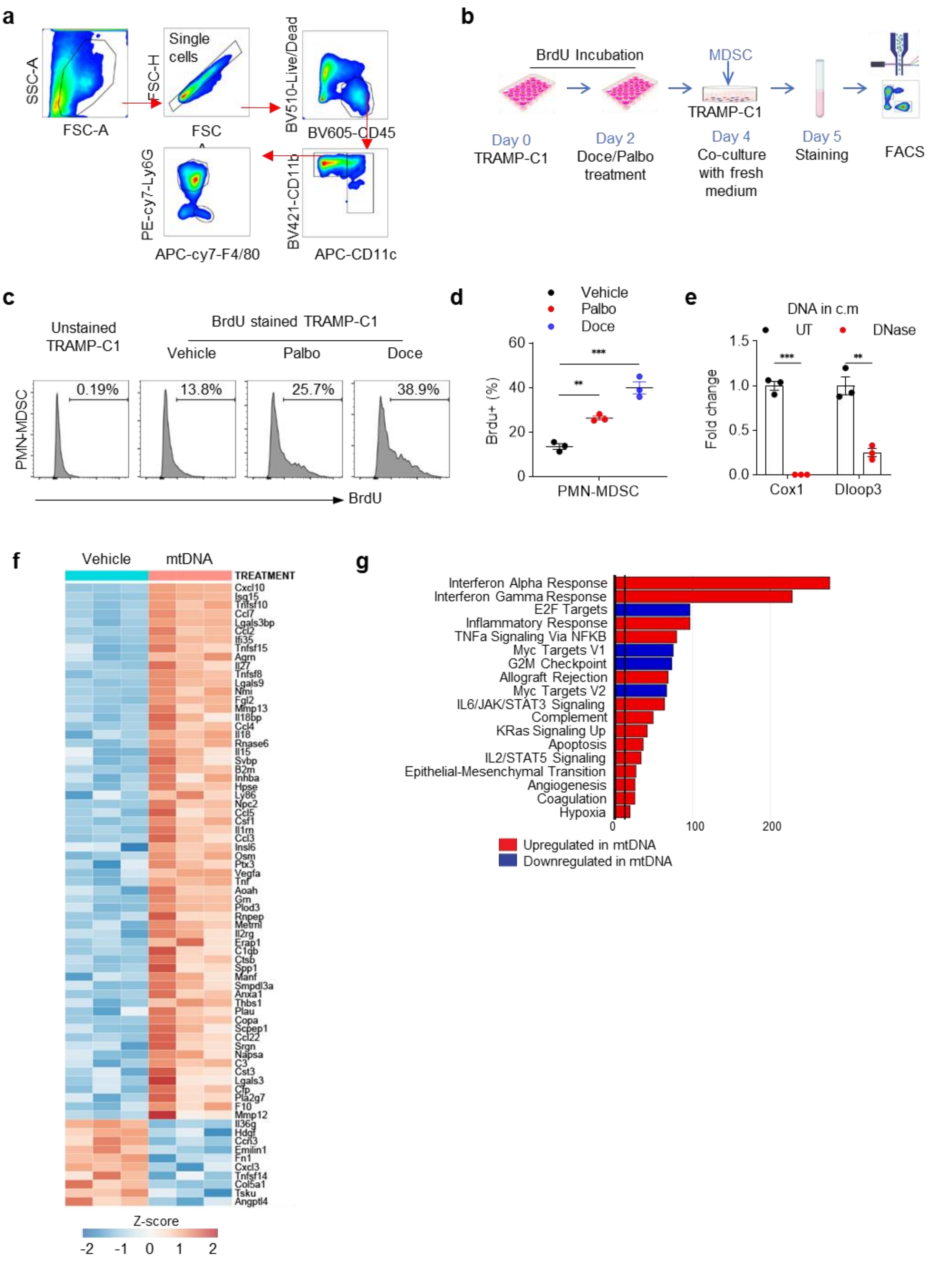


**Extended Data Fig. 7 PMN-MDSCs uptake mtDNA released by senescent tumor cells**

**a**, Gating strategy for mouse CD11b$^{+}$CD11c$^{+}$; CD11b$^{+}$F4/80$^{+}$ and CD11b$^{+}$Ly6G$^{+}$ myeloid cells from tumor xenograft in NRG mice. **b**, Scheme of timeline and experimental design of MDSC co-cultured with BrdU labeled TRAMP-C1 cells after Pablo and Doce treatment. **c**, **d**, Representative histogram (**c**) and quantification (**d**) of BrdU signal fluorescence intensity in PMN-MDSC ($CD11b^{+}$ $Ly6C^{Int}$ $Ly6G^{+}$) after co-cultured with TRAMP-C1 treated with Palbo or Doce. **e**, Quantification of mtDNA in c.m from Palbo-treated TRAMP-C1 cells performed with DNase treatments. **f**, Heat map of RNA-seq of BM-MDSC transfected with or without mtDNA. The color key represents the normalized Z score. **g**, Pathways/function changed in BM-MDSC transfected with mtDNA vs. Vehicle; Only pathways that were different between groups with $p < 0.01$ adjusted for multiple comparisons are shown. All values are presented as the mean ± SEM. One-way ANOVA followed by Tukey's multiple comparisons test was used in **d**, **e**. *$p < 0.05$; **$p < 0.01$; ***$p < 0.001$; ns, not significant.

### Extended Data Fig. 8

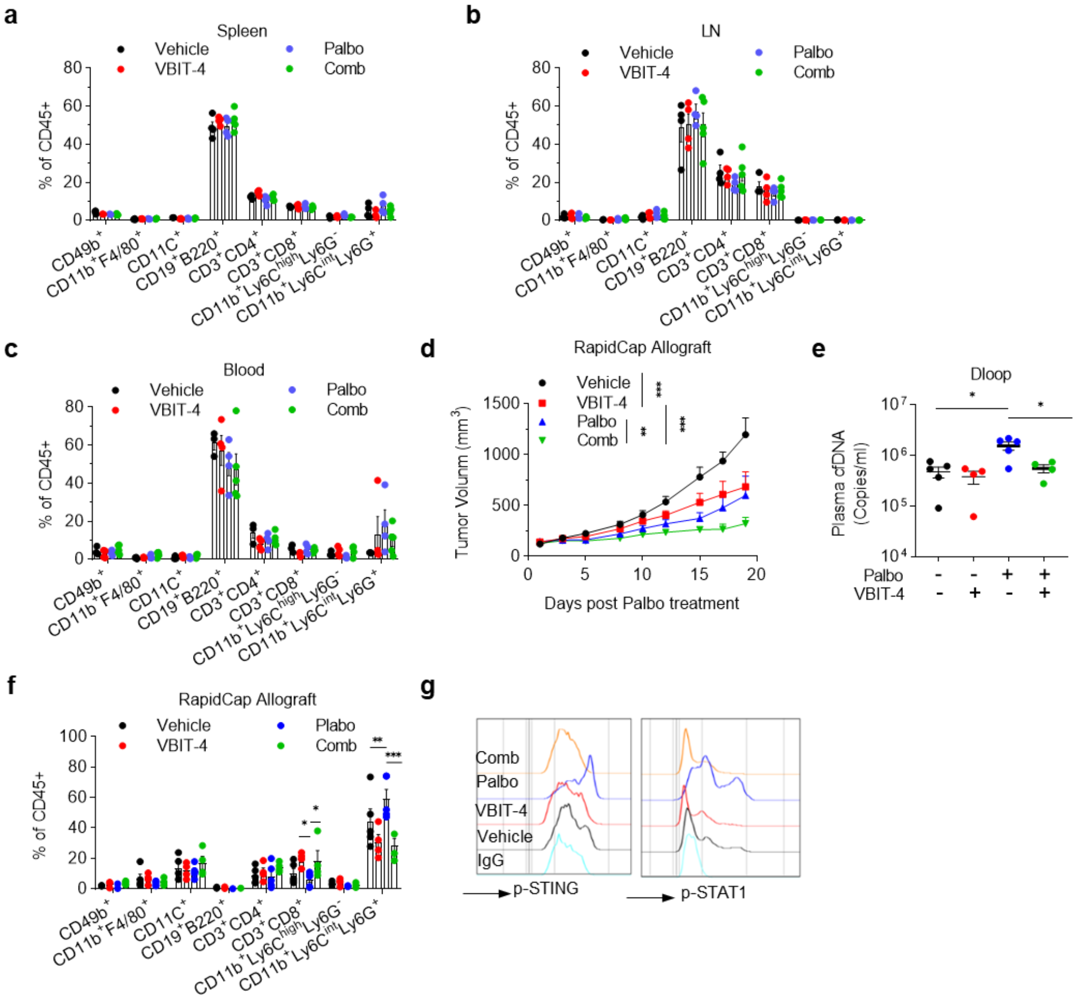


**Extended Data Fig. 8 VIBT4 treatment enhances TIS efficacy and decreases cGAS-STING activation in PMN-MDSCs**

**a-c**, Percentages of immune cell populations infiltration in the spleen (**a**), LN (**b**) and blood (**c**) (gated on $CD45^+$ cells). **d**, RapidCap tumor growth in mice with Palbo, VBIT-4, or combination treatment. UT: n =9, VBIT-4: n=6, Palbo: n=7, Comb: n=7. **g**, Representative histogram of p-STING and p-STAT1 in $CD11b^+Ly6C^{int}Ly6G^+$ PMN-MDSC sorted from TRAMP-C1 tumors with Palbo, VBIT-4, or combination treatment. All values are presented as the mean ± SEM. Two-way ANOVA followed by Dunnett's multiple comparisons test was used to evaluate the statistical significance in **a-f**. *p < 0.05; **p < 0.01; ***p < 0.001; NS, not significant.
